# VHL synthetic lethality screens uncover CBF-β as a negative regulator of STING

**DOI:** 10.1101/2024.09.03.610968

**Authors:** James A C Bertlin, Tekle Pauzaite, Qian Liang, Niek Wit, James C Williamson, Jia Jhing Sia, Nicholas J Matheson, Brian M Ortmann, Thomas J Mitchell, Anneliese O Speak, Qing Zhang, James A Nathan

## Abstract

Clear cell renal cell carcinoma (ccRCC) represents the most common form of kidney cancer and is typified by biallelic inactivation of the von Hippel-Lindau (*VHL*) tumour suppressor gene. Here, we undertake genome-wide CRISPR/Cas9 screening to reveal synthetic lethal interactors of *VHL*, and uncover that loss of Core Binding Factor β (CBF-β) causes cell death in *VHL*-null ccRCC cell lines and impairs tumour establishment and growth *in vivo*. This synthetic relationship is independent of the elevated activity of hypoxia inducible factors (HIFs) in *VHL*-null cells, but does involve the RUNX transcription factors that are known binding partners of CBF-β. Mechanistically, CBF-β loss leads to upregulation of type I interferon signalling, and we uncover a direct inhibitory role for CBF-β at the *STING* locus controlling Interferon Stimulated Gene expression. Targeting CBF-β in kidney cancer both selectively induces tumour cell lethality and promotes activation of type I interferon signalling.

## Introduction

Kidney cancer is a major and growing cause of morbidity and mortality worldwide, with approximately 180,000 patients dying each year from the disease^1,2^. Clear cell renal cell carcinoma (ccRCC) is the most common histological subtype in adults, accounting for approximately 85% of cases^3,4^. While low grade tumours can be cured by partial or radical nephrectomy, the systemic medical options to manage metastatic ccRCC remain largely inadequate, and only 13% of patients with metastatic spread at the time of diagnosis survive 5 years^5^.

Biallelic inactivation of the *VHL* tumour suppressor gene at chromosome 3p25 is a hallmark event in the development of ccRCC tumours^6,7^. Loss-of-heterozygosity of chromosome 3p occurs in childhood or adolescence, followed decades later by mutation or epigenetic inactivation of the remaining *VHL* allele. The subsequent acquisition of subclonal driver mutations, including the chromatin regulators *PBRM1*, *SETD2* and *BAP1*, culminates in the establishment of overt ccRCC tumours^8–10^. Inherited *VHL* mutations also underlie the hereditary autosomal dominant von Hippel-Lindau disease, in which patients develop numerous neoplasms early in life^11^.

VHL forms the substrate recognition component of the E3 ligase complex responsible for the ubiquitylation of prolyl-hydroxylated hypoxia inducible factor (HIF)-α subunits under conditions of abundant molecular oxygen^12,13^. In hypoxic environments, or upon genetic depletion of VHL, HIF-α accumulates, heterodimerises with HIF-1β/ARNT, and translocates to the nucleus, whereupon it drives a diverse transcriptional program, including genes implicated in angiogenesis and glycolytic metabolism^14^. The stabilisation of the HIF-2α isoform in particular accounts for many of the distinctive features of ccRCCs: vascular endothelial growth factor (VEGF) secretion results in highly angiogenic tumours, while increased fatty acid biosynthesis and impaired mitochondrial fatty acid transport causes cytoplasmic lipid deposition and the characteristic clear cell morphology^15–18^.

The term ‘synthetic lethality’ is used to describe the phenomenon in which the perturbation of two individually non-essential genes is lethal to a cell^19^. This paradigm has been used to target tumours with characteristic mutational drivers, exemplified by the employment of PARP inhibitors in the treatment of BRCA1/2-defective breast and ovarian cancers^20–23^. Over the last two decades, several groups have employed genetic and chemical screening platforms, computational algorithms, and hypothesis-driven approaches to identify synthetic lethal interactors of *VHL*^24–34^. However, while some hits may hold potential for the specific treatment of ccRCC, there has been little concordance between studies, presumably reflecting the diversity of cell lines used and the limitations of existing approaches, including the off-target effects of RNA interference techniques and the use of sub-optimal libraries.

Utilising pooled genome-wide CRISPR/Cas9 mutagenesis in ccRCC cell lines, we uncover synthetic lethal interactions for *VHL*, and find that Core Binding Factor β (CBF-β) loss selectively results in decreased tumour growth in ccRCCs. Mechanistically, CBF-β deficiency promotes cell death in ccRCCs, but we also find an unanticipated role for CBF-β in the repression of type I interferon (IFN) signalling. Loss of CBF-β results in Interferon Stimulated Gene (ISG) expression, resulting from the de-repression of STING (Stimulator of Interferon Genes) transcription. CBF-β therefore tunes cell-intrinsic type I interferon signalling, and its loss sensitises cells to a heightened STING-mediated immune response. Therefore, we propose that CBF-β offers a potential therapeutic target in ccRCC through its synthetic lethal relationship with *VHL* loss.

## Results

### Genome-wide CRISPR/Cas9 screening reveals synthetic viability interactors of *VHL*

To find genes that altered the viability of *VHL*-null cells, we employed the widely-used ccRCC models 786O and RCC4 alongside paired VHL-reconstituted lines (786O+*VHL* and RCC4+*VHL*) (**Fig. 1a and Extended Data** Fig. 1a)^12,35,36^. Pooled CRISPR/Cas9 screening was performed by transducing isogenic cell lines expressing Cas9 with the Toronto KnockOut version 3.0 (TKOv3) genome-wide sgRNA library, passaging cells until 17 population doublings had occurred, and undertaking deep sequencing of sgRNAs (**Fig. 1b and Extended Data** Fig. 1b)^37^. Identification of a curated set of core essential genes that dropped out over the duration of the culture confirmed the efficiency of the screen (**Extended Data** Fig. 1c **and Supplementary Table 1**)^38^. Synthetic viability interactors of *VHL* were identified as those targeted by sgRNA sequences which were relatively less or more abundant in the *VHL*-null lines at the conclusion of the screen (**Fig. 1c and Supplementary Table 1**). The positive controls *ARNT*/*HIF1β* and *MTOR* dropped out in the *VHL*-null setting, confirming the validity of our approach^34,39^. Alongside these expected findings we identified a putative synthetic lethal relationship between *VHL* and *CBFB* (encoding CBF-β) in both ccRCC backgrounds (**Fig. 1c,d**). CBF-β is best characterised as forming a transcriptional complex with RUNX proteins to activate or repress target genes^40,41^. Both *RUNX1* and *RUNX2* were identified in the RCC4 screen (**Fig. 1c**), consistent with a potential role of these transcriptional complexes in the synthetic viability relationship with *VHL*.

**Fig. 1.**
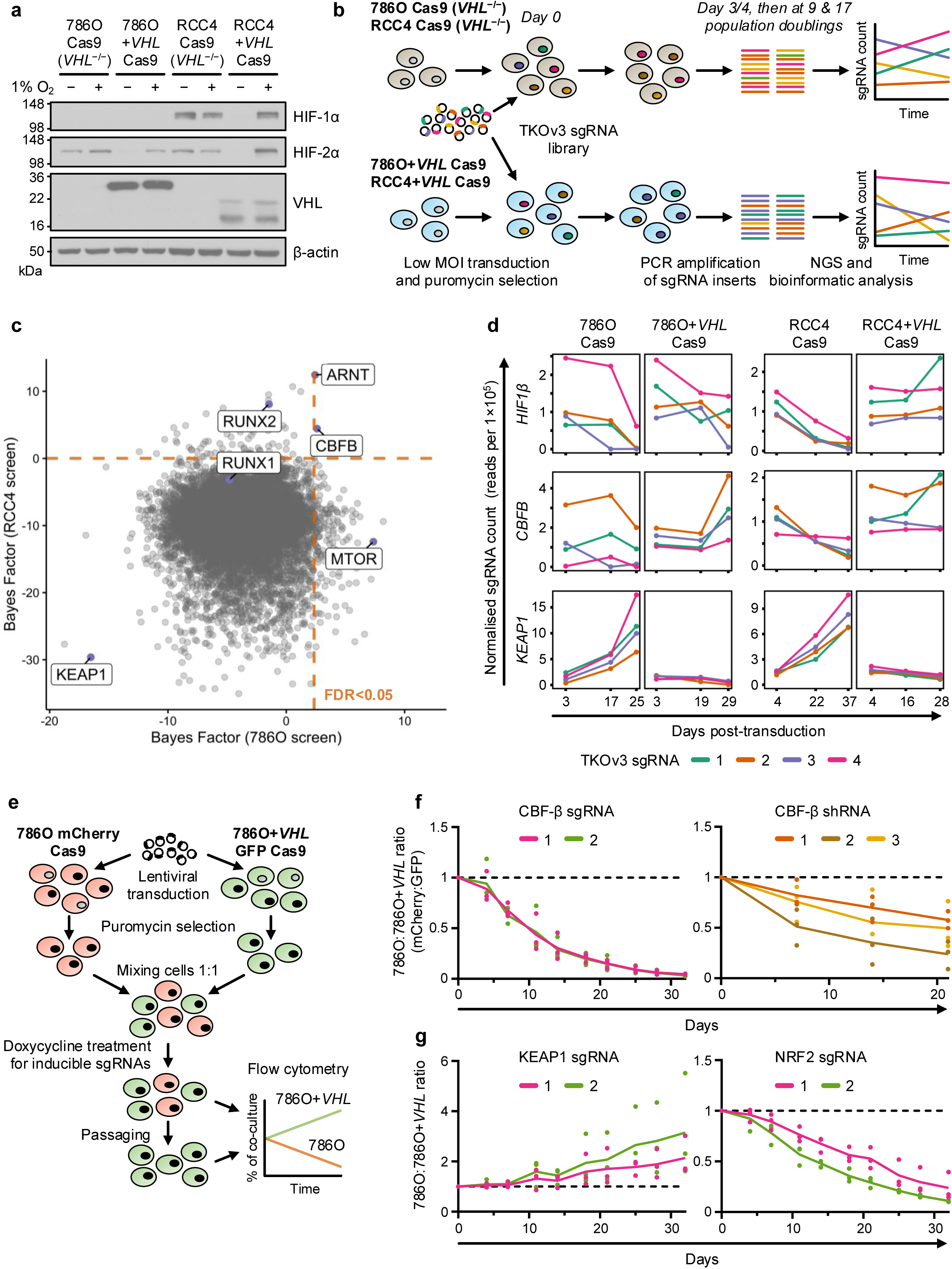
Genome-wide CRISPR/Cas9 screening reveals synthetic viability interactors of *VHL* (a) HIF-α levels in ccRCC cells and paired *VHL*-reconstituted cells after culture at either 1% or 21% O_2_ for 24 hours. Immunoblot representative of 3 independent experiments. (b) Schematic of CRISPR/Cas9 screen design. Paired *VHL*-null and -reconstituted cell lines were transduced with the genome-wide TKOv3 sgRNA library. Cells were passaged in parallel for several weeks, and genomic DNA extracted at early, intermediate and late timepoints for the identification of sgRNA abundance by Next Generation Sequencing (NGS). sgRNAs targeting synthetic lethal interactors of *VHL* are selectively depleted in the *VHL*-null 786O Cas9 and RCC4 Cas9 backgrounds. (c) Pairwise analysis of the late timepoints of 786O Cas9 vs. 786O+*VHL* Cas9 cells (x-axis) and RCC4 Cas9 vs. RCC4+*VHL* Cas9 cells (y-axis) from CRISPR/Cas9 screens. A higher Bayes Factor calculated by BAGEL2 indicates a more robust synthetic lethal relationship between the given gene and *VHL*. Orange lines denote FDR<0.05. (d) Normalised sgRNA counts from CRISPR/Cas9 screens. (e) Competitive growth assay method. Transduced fluorophore-expressing 786O and 786O+*VHL* cells are mixed 1:1, passaged and harvested for flow cytometry analysis. 100 ng/ml doxycycline is added to experiments involving inducible sgRNAs to initiate gene editing. (**f,g**) Validation of functional interactions between *VHL* and *CBFB* (**f**), or between *VHL* and *KEAP1* and *NRF2* (**g**), by competitive growth assay using doxycycline-inducible sgRNAs or constitutively-expressed shRNAs. n=3 biologically independent replicates. One-way ANOVA based on Area Under the Curve compared to empty vector or scrambled control: CBF-β sgRNA 1 P<0.0001; CBF-β sgRNA 2 P<0.0001; CBF-β shRNA 1 P=0.14; CBF-β shRNA 2 p=0.0034; CBF-β shRNA 3 P=0.042; KEAP1 sgRNA 1 P=0.42; KEAP1 sgRNA 2 P=0.11; NRF2 sgRNA 1 P=0.0002; NRF2 sgRNA 2 P<0.0001.

*KEAP1* was identified as the top hit that when mutagenized conferred a proliferative advantage to *VHL*-null cells (**Fig. 1c,d**). KEAP1 ubiquitylates and traps NRF2 in the cytoplasm to prevent NRF2-mediated transcription of antioxidant factors including *NQO1* and *HMOX1*^42,43^, raising the possibility that upregulation of the NRF2 response may confer a survival advantage following *VHL* loss.

We developed a competitive growth assay with a fluorescent read-out to validate target genes (**Fig. 1e**). By labelling the 786O *VHL*-null cells with mCherry and the *VHL*-reconstituted cells with GFP, we could readily distinguish a growth advantage or disadvantage when both cell lines were mixed in a 1:1 ratio (**Fig. 1e**). Inducible depletion of CBF-β, using both sgRNA and shRNA, confirmed the *VHL*/*CBFB* synthetic lethal relationship (**Fig. 1f and Extended Data** Fig. 1d,e). Conversely, KEAP1 loss increased the growth of 786O cells relative to 786O+*VHL* cells (**Fig. 1g and Extended Data** Fig. 1f). We also found that NRF2 was synthetic lethal with VHL in the 786O competitive growth assay (**Fig. 1g**), but did not alter the levels of HIF-2α or HIF target gene expression (**Extended Data** Fig. 1f,g). Therefore, the KEAP1-NRF2 axis may protect against a greater degree of oxidative stress in VHL-deficient cells.

### Deletion of *CBFB* causes cell death in *VHL*-null cells

We chose to focus on *CBFB* given the potential synthetic lethal relationship between *CBFB* and *VHL*, the additional identification of *RUNX* genes in the RCC4 screen, and the observation that high expression of *CBFB* mRNA in ccRCC tumours correlated strongly with poor survival outcomes in ccRCC tumours in The Cancer Genome Atlas database (**Fig. 2a**).

**Fig. 2.**
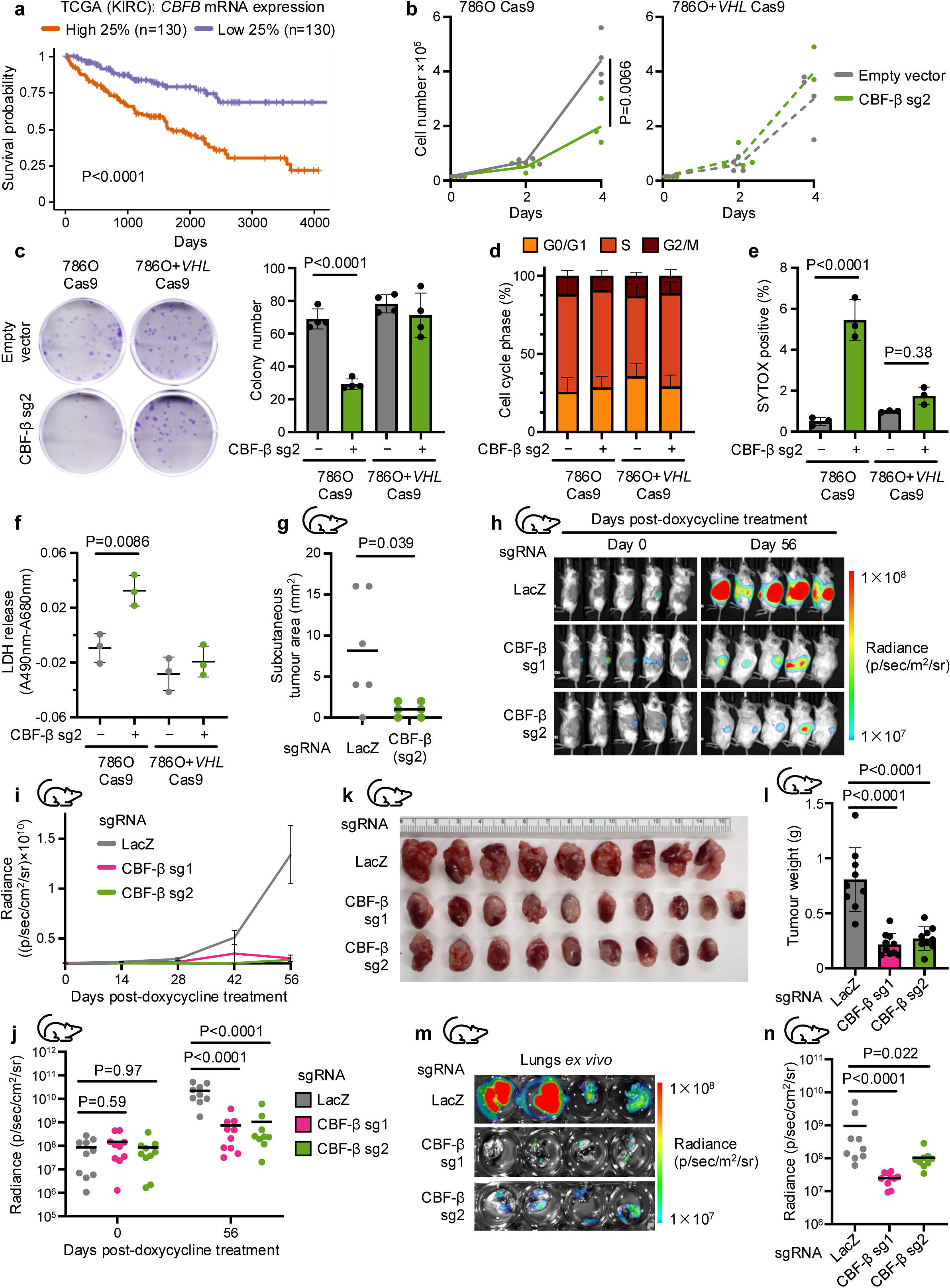
CBF-β loss causes cell death in VHL-null ccRCCs and impairs tumour growth *in vivo* (a) High expression of *CBFB* in kidney cancer is associated with poor outcomes. Kaplan-Meier survival analysis of TCGA data for ccRCC^7^, comparing tumours in the highest and lowest quartiles of *CBFB* mRNA expression. n=130 patients for each group. Log-rank test. (b) Proliferation assay of 786O Cas9 and 786O+*VHL* Cas9 cells transduced with an sgRNA targeting CBF-β (CBF-β sg2), or an empty vector control. n=4 biologically independent replicates. Two-way ANOVA of cell number at day 4. (c) Cells were transduced with CBF-β sg2 or an empty vector control and analysed by clonogenic assay. Representative image of colonies after staining with 0.5% crystal violet solution. n=4 biologically independent replicates. Mean ± SD. Two-way ANOVA. (d) Proportion of cells in G0/G1, S, and G2/M cell cycle phases in asynchronous 786O Cas9 and 786O+*VHL* Cas9 populations transduced with CBF-β sg2 or an empty vector control, determined by BrdU incorporation and propidium iodide staining. n=4 biologically independent replicates. Mean ± SD. (e) SYTOX AADvanced cell death assay, indicating the proportion of 786O Cas9 and 786O+*VHL* Cas9 cells stained positive following transduction with CBF-β sg2 or an empty vector control. n=3 biologically independent replicates. Mean ± SD. Two-way ANOVA. (f) LDH cytotoxicity assay, quantifying the amount of LDH in culture supernatant in 786O Cas9 and 786O+*VHL* Cas9 cells transduced with CBF-β sg2 or an empty vector control. Background absorbance at 680 nm was subtracted from the 490 nm signal, and readings normalised to a cell-free unconditioned media control. n=3 biologically independent replicates. Mean ± SD. Two-way ANOVA. (g) CBF-β is required for tumour establishment *in vivo*. Tumour area 28 days after subcutaneous injection of 786O Cas9 cells transduced with sgRNAs targeting CBF-β or LacZ into NSG mice. n=6 mice per group. Mann-Whitney U test. (**h**-**n**) Xenograft model of luciferase-expressing 786O Cas9 cells injected orthotopically into the left kidneys of NSG mice, using doxycycline treatment to induce the expression of sgRNAs targeting CBF-β (sg1 and sg2) or LacZ as a control. Representative bioluminescence imaging of mice at day 0 and day 56 after the initiation of doxycycline treatment (**h**). Quantification of the bioluminescence of primary tumours following doxycycline treatment (**i,j**). Left kidneys following euthanasia at day 56 post-doxycycline treatment, with tumour mass quantified by the subtraction of right kidney mass from left kidney mass (**k,l**). Representative and quantified bioluminescence of isolated lungs harvested from mice at day 56 post-doxycycline treatment (**m,n**). n=10 mice per group, of which 1 died between day 42 and day 56 post-treatment in each of the LacZ and CBF-β sg2 groups. Mean ± SEM (**i**) or SD (**l**). One-way ANOVA.

We first sought to characterise the proliferation phenotype of CBF-β loss in ccRCC cells. The growth and colony formation potential of 786O cells was significantly impaired by CBF-β depletion, but this growth defect was abrogated in the isogenic *VHL*-reconstituted line (**Fig. 2b,c**). CBF-β loss also impeded growth in some other *VHL*-null ccRCC lines (A498 and RCC10 cells), but not in 769P ccRCC cells, or the HKC8 model of healthy proximal tubule epithelium (**Extended Data** Fig. 2a,b). Prior studies report differing sensitivities to HIF-2α inhibitors in ccRCC lines, and we considered whether an interaction with CBF-β and HIF-2α could explain the varied sensitivity of ccRCC lines to impaired growth with *CBFB* loss^39^. However, the functional interaction between *CBFB* and *VHL* was independent of HIF signalling, as the same phenotype remained in 786O cells following clonal knockout of HIF1β (**Extended Data** Fig. 2a,b, final columns), and *CBFB* knockout did not consistently affect the levels or transcriptional activity of HIF-2α (**Extended Data** Fig. 2c,d).

The proliferation defect in 786O cells could be caused by either a defect in cell cycling, or an increased rate of cell death. We excluded a significant deficiency in cell cycling as neither the distribution of cells within cycle phases nor the progression of synchronised populations through the cell cycle were affected by *CBFB* knockout (**Fig. 2d and Extended Data** Fig. 2e). Rather, the depletion of CBF-β caused cytotoxicity in *VHL*-null cells, demonstrated by SYTOX AADvanced staining and quantification of released lactate dehydrogenase (LDH) (**Fig. 2e,f**). We investigated the mode of cell death using assays of apoptosis, or specific inhibitors of necroptosis (Nec-1s), pyroptosis (VRT-043198), and ferroptosis (ferrostatin). Apoptosis assays did not demonstrate a clear induction of this pathway following CBF-β depletion (**Extended Data** Fig. 2f,g), and treatment with the highest tolerable doses of the various cell death inhibitors did not restore growth in the CBF-β-deficient 786O cells (**Extended Data** Fig. 2h). Therefore, while the precise modality of cell death is unclear, our studies indicate that CBF-β loss does lead to cytotoxicity in 786O cells.

To establish whether CBF-β loss also impaired tumourigenesis *in vivo*, we performed xenograft experiments in immunodeficient NSG mice. First, we observed that CBF-β knockout severely restricted the ability of 786O cells to develop tumours in the subcutaneous microenvironment (**Fig. 2g**). Next, to interrogate the more clinically-relevant scenario of established kidney tumours, we employed an orthotopic model of ccRCC using luciferase-expressing 786O cells harbouring doxycycline-inducible sgRNA vectors^44^. Upon confirmation of tumour growth in each kidney, as indicated by increased luciferase activity over time, we administered doxycycline chow to mice in order to induce CBF-β depletion *in vivo*. Deletion of CBF-β with two independent sgRNAs both substantially impeded tumour growth (**Fig. 2h-l**) and prevented metastasis to the lungs (**Fig. 2m,n**), confirming the relevance of the CBF-β/VHL synthetic lethality *in vivo*.

### CBF-β and RUNX proteins are required for the synthetic lethal interaction with VHL

Given the canonical transcriptional function of CBF-β in complex with RUNX1-3, and the manifestation of RUNX2 as a synthetic lethal hit in the RCC4 CRISPR screen (**Fig. 1c**), we hypothesised that *CBFB* loss could result in the death of *VHL*-null cells via impaired RUNX activity. Indeed, the combined knockout of RUNX1 and RUNX2, but not of either isoform individually, caused a similar degree of synthetic lethality to CBF-β loss in a competitive growth assay (**Fig. 3a**), and the deletion of all three RUNX proteins mimicked the observed CBF-β proliferation phenotype in 786O cells (**Fig. 3b**). However, the synthetic lethal effect of CBF-β with VHL loss could not be simply explained by a reduction of complex formation with RUNX proteins, as we noted that deletion of CBF-β caused a profound decline in RUNX1 and RUNX2 protein abundance in both 786O and 786O+*VHL* cells (**Fig. 3c,d**).

**Fig. 3.**
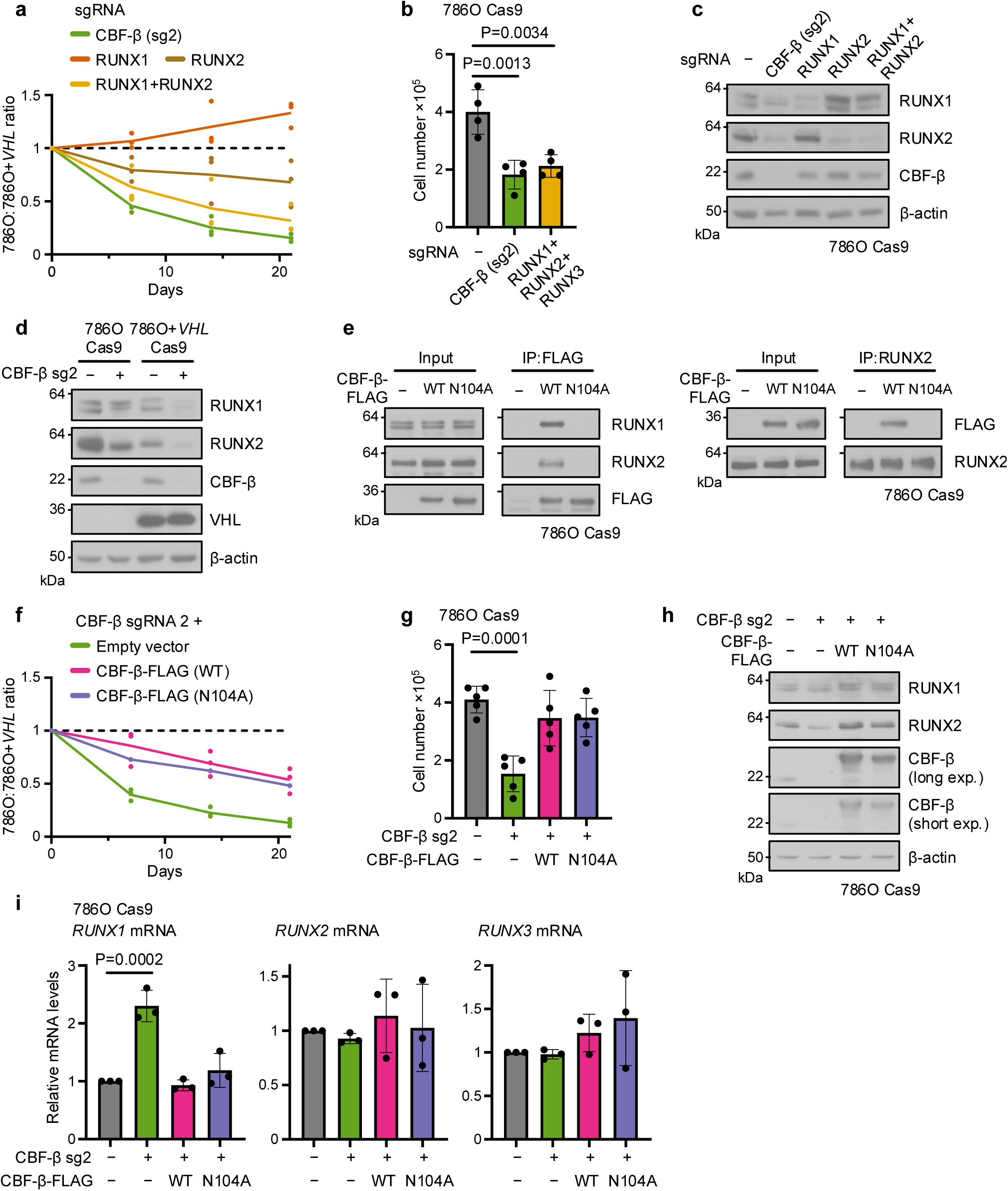
Post-transcriptional regulation of RUNX1 and RUNX2 by CBF-β (a) Combined loss of *RUNX1* and *RUNX2* functionally mimics *CBFB* deletion. Competitive growth assay of cells transduced with sgRNAs targeting CBF-β, RUNX1, RUNX2, or both RUNX1 and RUNX2. n=3 biologically independent replicates. One-way ANOVA based on Area Under the Curve compared to empty vector control. CBF-β sgRNA 2 P=0.0003; RUNX1 sgRNA P=0.31; RUNX2 sgRNA P=0.12; RUNX1 sgRNA + RUNX2 sgRNA P=0.0024. (b) Proliferation assay of 786O Cas9 cells transduced with sgRNAs targeting CBF-β or RUNX1, RUNX2 and RUNX3 in combination, or an empty vector control. n=4 biologically independent replicates. Mean ± SD. One-way ANOVA. (c) CBF-β loss depletes RUNX protein levels. Immunoblot of cells transduced with sgRNAs targeting CBF-β, RUNX1, RUNX2, or both RUNX1 and RUNX2. Representative of 3 biologically independent replicates. (d) Immunoblot of 786O Cas9 and 786O+*VHL* Cas9 cells transduced with CBF-β sg2 or an empty vector control. Representative of 3 biologically independent replicates. (e) Cells were transduced with overexpression vectors encoding CBF-β-FLAG (WT), CBF-β- FLAG (N104A), or an empty vector. FLAG-tagged CBF-β and endogenous RUNX2 were immunoprecipitated (IP) from cell lysates as indicated. Immunoblots representative of 3 biologically independent replicates. (f) RUNX-binding mutant of CBF-β restores 786O Cas9 cell growth to the same level as wild- type (WT) CBF-β following *CBFB* knockout. Competitive growth assay of cells transduced with CBF-β sg2 and overexpression vectors encoding CBF-β-FLAG (WT), CBF-β-FLAG (N104A), or an empty vector control. n=3 biologically independent replicates. One-way ANOVA based on Area Under the Curve compared to empty sgRNA vector control: CBF-β sgRNA 2 + empty vector: P=0.0002; CBF-β sgRNA 2 + CBF-β-FLAG (WT) P=0.066; CBF-β sgRNA 2 + CBF-β-FLAG (N104A) P=0.017. One-way ANOVA based on Area Under the Curve compared to CBF- β sgRNA 2: CBF-β sgRNA 2 + CBF-β-FLAG (WT) P=0.0048; CBF-β sgRNA 2 + CBF-β-FLAG (N104A) P=0.017. (g) Proliferation assay of 786O Cas9 cells transduced with CBF-β sg2 and overexpression vectors encoding CBF-β-FLAG (WT) or CBF-β-FLAG (N104A), or an empty vector control. n=4 biologically independent replicates. Mean ± SD. One-way ANOVA. (**h,i**) Regulation of RUNX by CBF-β is post-transcriptional. 786O Cas9 cells were transduced with CBF-β sg2 and overexpression vectors encoding CBF-β-FLAG (WT) or CBF-β-FLAG (N104A), or an empty vector control. RUNX protein and mRNA levels were analysed by immunoblot (**h**) and qPCR (**i**). n=3 biologically independent replicates. Mean ± SD. One-way ANOVA.

To further interrogate the role of CBF-β-RUNX dimerisation, we expressed FLAG-tagged overexpression constructs of wild-type (WT) CBF-β and an N104A-mutant form, which fails to bind the Runt homology domain of RUNX proteins^45^, in 786O or CBF-β-deficient 786O cells. We confirmed that the N104A mutation prevented RUNX binding by co-immunoprecipitation (**Fig. 3e**). Importantly, the sgRNA used to induce CBF-β deletion (CBF-β sg2) was directed against the boundary between intron 2 and exon 3 of the *CBFB* locus and therefore permitted knockout of the endogenous CBF-β protein while sparing the overexpression constructs. However, overexpression of either wild-type or N104A CBF-β restored the proliferation of CBF- β-null 786O cells to a similar extent (**Fig. 3f,g**). Moreover, we noted both CBF-β constructs restore RUNX protein levels without increasing *RUNX1-3* mRNA expression (**Fig. 3h,i**). Together, these findings indicate that while RUNX proteins are required for the *CBFB* synthetic lethal growth defect in 786O cells, the direct association of CBF-β with RUNX proteins may not be necessary.

### *CBFB* loss induces a cell intrinsic interferon response

As CBF-β and RUNX appeared to exercise their functions at both transcriptional and translational levels, we undertook RNA sequencing and tandem mass tag (TMT)-labelled mass spectrometry to characterise the effects of CBF-β loss in VHL-proficient and -deficient backgrounds (**Fig. 4a**). Strikingly, in 786O cells, *CBFB* knockout caused a net increase in transcription and specifically induced the expression of a plethora of ISGs exclusively in *VHL*- null 786O cells (**Fig. 4b-d and Supplementary Table 2, 3**). Quantitative PCR (qPCR) analysis confirmed that the upregulation of several ISGs (*IFIT1*, *OASL*, *ISG15*, and *RSAD2*) was specific to CBF-β depletion, using both sgRNA and shRNA (**Extended Data** Fig. 3a-c), and prevented by reconstitution with both wild-type and N104A-mutant CBF-β-FLAG constructs (**Fig. 4e and Extended Data** Fig. 3d). The selective ISG stimulation in *VHL*-null cells was also independent of HIF activity, as it was not abolished by *HIF1β* knockout (**Fig. 4f**), and observed in other but not all ccRCC lines (**Extended Data** Fig. 3e).

**Fig. 4.**
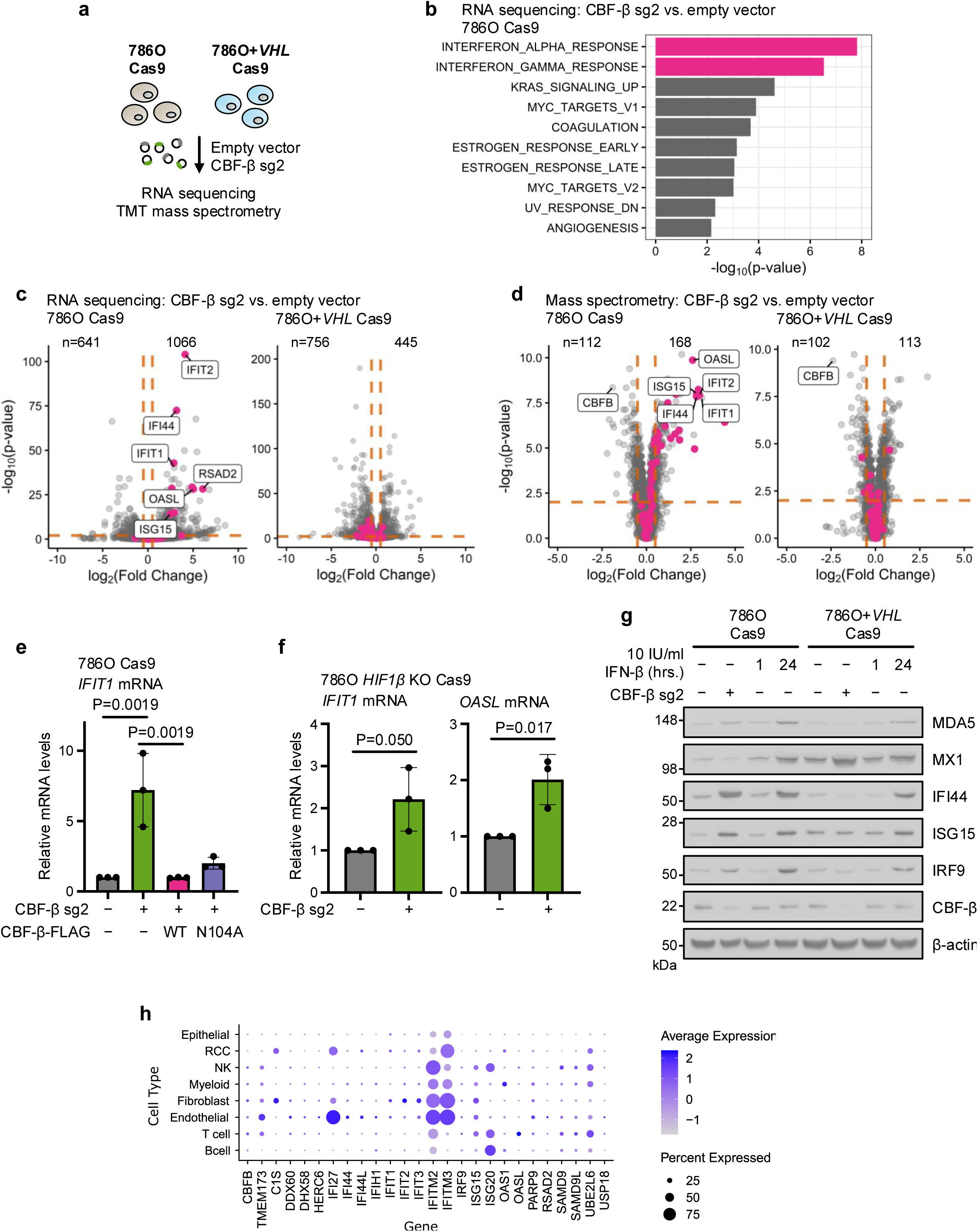
CBF-β loss induces an Interferon Stimulated Gene (ISG) signature (a) 786O Cas9 and 786O+*VHL* Cas9 cells were analysed by RNA sequencing and tandem mass tag (TMT)-labelled LC-MS to assess global changes in mRNA and protein upon CBF-β depletion relative to empty vector-transduced cells. (b) Gene Set Enrichment Analysis of Hallmark gene sets from RNA sequencing analysis of differential gene expression in 786O Cas9 cells following *CBFB* knockout^100^. n=3 biologically independent replicates. (**c,d**) Volcano plots of RNA sequencing (**c**) and mass spectrometry (**d**) data in the conditions outlined in (**a**). Orange lines indicate log_2_(Fold Change) ± 0.5, and P_adj_=0.01. Pink dots highlight genes represented within the Hallmark INTERFERON_ALPHA_RESPONSE. Number of genes upregulated or downregulated (determined by the log2(Fold Change) and Padj cut-off values above) are specified. Data points represent the mean value for each gene from 3 biologically independent replicates. (e) *IFIT1* is specifically upregulated by CBF-β loss. qPCR analysis of *IFIT1* expression in 786O Cas9 cells upon transduction with CBF-β sg2 and overexpression vectors encoding CBF-β- FLAG (WT) or CBF-β-FLAG (N104A), or an empty vector control. n=3 biologically independent replicates. Mean ± SD. One-way ANOVA. (f) qPCR analysis of ISGs in clonal 786O HIF1β knockout Cas9 cells upon CBF-β sg2 transduction, relative to empty vector-transduced controls. n=3 biologically independent replicates. Mean ± SD. Unpaired *t*-test. (g) Immunoblot of ISGs in 786O Cas9 and 786O+*VHL* Cas9 cells transduced with CBF-β sg2 or an empty vector control. Controls were treated with 10 IU/ml IFN-β for 1 or 24 hours. Representative of 3 biologically independent replicates. (h) Analysis of single cell transcriptomic data from patients with ccRCC. The average expression and the percentage of cells that express type I interferon genes is shown for the principal cell types within the tumour micro-environment.

We next determined how the induction of ISGs in CBF-β-deficient 786O and 786O+*VHL* cells compared with type I IFN treatment by titrating levels of IFN-β. We observed that CBF-β loss was equivalent to treatment with approximately 24 hours of 10 IU/ml IFN-β (**Fig. 4g**). CBF-β loss also induced ISGs non-uniformly, affecting a subset of IFN-β-responsive genes (IFI44, ISG15, and IRF9), but not MDA5 or MX1 (**Fig. 4g**). Given this low amplitude ISG response, we considered whether the induction of ISGs related to lentiviral transduction of our sgRNA targeting *CBFB*. However, ISGs were not induced if cells were transduced with sgRNAs of similar G:C content targeting alternative loci to *CBFB* (**Extended Data** Fig. 3f). Thus, CBF-β loss promotes an ISG response in 786O cells.

To further explore ISG expression in VHL-deficient renal cancer, we analysed the single cell transcriptional profiles of treatment-naive renal tumours from 12 patients, who underwent surgical resection^46^. Using the same type I IFN gene signatures as for the bulk RNA-seq (**Fig. 4b,c**) we observed that there was considerable heterogeneity in ISG expression across the different cell populations within the tumour (**Fig. 4h, Supplementary** Fig. 1). RCC cells showed only low levels of ISGs, with the bulk of ISG expression stemming from the endothelial cells and some immune subsets (**Fig. 4h, Supplementary** Fig. 1). *CBFB* transcript levels were low across all cell types, therefore correlating *CBFB* expression with an ISG expression was not possible. However, these transcriptional profiles are consistent with general repression of ISGs within the tumour cells.

### A STING-TBK1-IRF3 axis drives ISG expression in VHL-deficient renal cancer cells

The finding that *CBFB* loss resulted in type I IFN expression suggested that CFB-β either negatively regulated IFN signalling or that it resulted in damage signals detected by pattern recognition receptors (PRRs). ISG expression is classically stimulated when microbial nucleic acids are sensed by PRRs which converge, via the adaptors STING, MAVS and TRIF, on the TBK1/IKK-ε kinases to promote the phosphorylation and dimerisation of IRF3^47–49^, activating ISGs and type I IFN expression (**Fig. 5a**). Upon secretion, type I IFNs then act in a paracrine or autocrine manner to initiate a JAK-STAT phosphorylation cascade which culminates in the transcription of ISGs by STAT1/2 heterodimers (in association with IRF9) or, less commonly, STAT1 homodimers (**Fig. 5a**)^50,51^. Our finding that *CBFB* deletion only altered the growth of 786O but not 786O+*VHL* cells in co-culture (**Fig. 1f**) suggested that type I IFN secretion was unlikely to explain the induction of ISGs, and that a cell intrinsic pathway was involved. We therefore anticipated that abolition of STAT1/2 signalling would abolish the IFN response to CBF-β loss. To our surprise, however, knockout of *STAT1*, *STAT2* or *IRF9* individually, or in combination, did not prevent *IFIT1* induction (**Fig. 5b and Extended Data** Fig. 4a-c), and we also did not observe phosphorylation of STAT1 at Tyr701 with *CBFB* knockout (**Fig. 4d**).

**Fig. 5.**
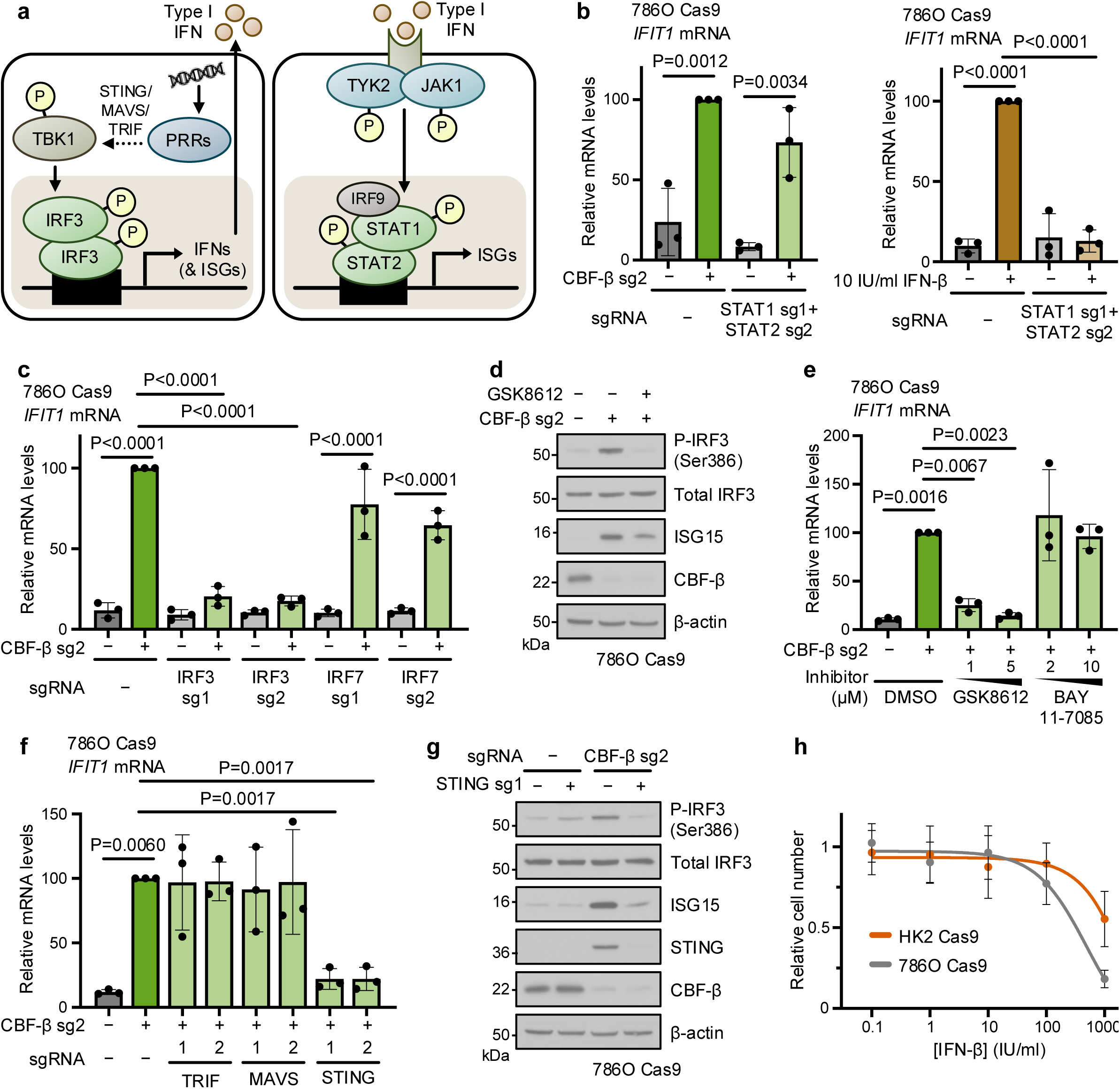
*CBFB* loss induces a STING-TBK1-IRF3 cell-intrinsic ISG response (a) Schematic of the type I IFN signalling pathway. Aberrant or foreign nucleic acids are detected by pattern recognition receptors (PRRs), which signal through adaptor proteins to stimulate TBK1-mediated phosphorylation of IRF3. IRF3 heterodimers translocate to the nucleus to trigger transcription of IFNs and a subset of ISGs. Secreted IFNs act through IFNAR receptors to promote STAT1- and STAT2-mediated ISG expression. (b) qPCR analysis of *IFIT1* expression in 786O Cas9 cells transduced with a vector encoding sgRNAs targeting both STAT1 (sg1) and STAT2 (sg2), or an empty vector control. Cells were additionally transduced with CBF-β sg2, or treated with 10 IU/ml IFN-β for 6 hours. n=3 biologically independent replicates. Mean ± SD. Two-way ANOVA. (c) ISG expression is dependent on IRF3. qPCR analysis of cells transduced with sgRNAs targeting CBF-β (sg2), IRF3 and IRF7, or an empty vector control. n=3 biologically independent replicates. Mean ± SD. Two-way ANOVA. (d) Cells were transduced with CBF-β sg2 or an empty vector control, and treated with 5 μM GSK8612 for 24 hours (to inhibit TBK1), as indicated. Immunoblot representative of 3 biologically independent replicates. (e) 786O Cas9 cells were transduced with CBF-β sg2 or an empty vector control, treated for 24 hours with the indicated doses of the TBK1 inhibitor GSK8612, the NF-κB inhibitor BAY 11- 7085, or the DMSO vehicle, and analysed by qPCR. n=3 biologically independent replicates. Mean ± SD. One-way ANOVA. (f) CBF-β loss specifically activates STING. qPCR analysis of cells transduced with CBF-β sg2 alone or in combination with sgRNAs targeting the PRR adaptors TRIF, MAVS and STING, compared to an empty vector-transduced control. n=3 biologically independent replicates. Mean ± SD. One-way ANOVA. (g) Immunoblot of 786O Cas9 cells transduced with sgRNAs targeting CBF-β (sg2) and STING (sg1), or an empty vector control. Representative of 3 biologically independent replicates. (h) Dose-response relationship between IFN-β treatment and cell proliferation over 72 hours in 786O Cas9 cells and proximal tubule HK2 Cas9 cells. n=6 biologically independent replicates. Mean ± SD.

We therefore examined if CBF-β loss acted upstream in ISG induction by depleting IRF3. *IFIT1* expression was profoundly sensitive to *IRF3* knockout in the CBF-β-null cells (**Fig. 5c and Extended Data** Fig. 4e). Moreover, the ISG response observed upon CBF-β knockout was entirely dependent on IRF3, and not on the closely related IRF7 (**Fig. 5c and Extended Data** Fig. 4e)^52^, and was associated with phosphorylation of IRF3 at Ser386 (**Fig. 5d and Extended Data** Fig. 4f), which is essential for IRF3 activation^53^. To confirm that IRF3 phosphorylation was required for subsequent ISG expression, we treated CBF-β-null 786O cells with the TBK1 inhibitor, GSK8612. This treatment prevented IRF3 Ser386 phosphorylation and the subsequent ISG induction in a dose-dependent manner (**Fig. 5d,e**). Lastly, we established that STING, but not MAVS or TRIF, was required for the induction of ISGs following *CBFB* knockout (**Fig. 5f,g and Extended Data** Fig. 4g), identifying a STING-TBK1-IRF3 axis that is activated in a CBF-β loss-dependent manner.

We next considered whether the activation of the STING-TBK1-IRF3 axis was sufficient to explain the synthetic lethal phenotype observed between *CBFB* and *VHL*, but STING or IRF3 depletion by themselves did not fully reverse the synthetic interaction (**Extended Data** Fig. 4h-j). However, IFN-β treatment did increase cell death in 786O cells compared to immortalised renal tubular epithelial cells (**Fig. 5h**). Therefore, while activation of the IRF3 axis cannot fully explain the synthetic lethal phenotype, ccRCCs are sensitive to type I IFN induced cell death.

### CBF-β represses STING to tune the type I interferon response

We were intrigued by the notion that CBF-β may be a negative regulator of STING. We noted that not only did CBF-β loss result in a STING-dependent induction of ISGs, but that protein levels of STING increased in CBF-β-null 786O cells (**Fig. 5g**). Therefore, we tested whether CBF-β depletion altered STING expression and the sensitivity of the cGAS-STING pathway to double stranded DNA (dsDNA) (**Fig. 6a**). Firstly, in 768O cells, we observed that *STING* transcription was dramatically increased upon *CBFB* knockout but reduced by its overexpression (**Fig. 6b**). Moreover, using herring testes DNA (HT-DNA), a long ‘non-self’ DNA molecule which enables specific interrogation of cGAS-STING function (**Fig. 6c**)^54,55^, we found a striking synergistic induction of ISGs following CBF-β loss: while HT-DNA alone provided only modest *IFIT1* upregulation, we observed a dramatic induction in response to combined HT-DNA transfection and CBF-β sg2 transduction (**Fig. 6d**). This was accompanied by increased expression of *IFNB1* (encoding IFN-β), despite its transcription not being affected by *CBFB* knockout alone. Conversely, overexpression of the wild-type CBF-β-FLAG construct blunted the response to HT-DNA (**Fig. 6d**).

**Fig. 6.**
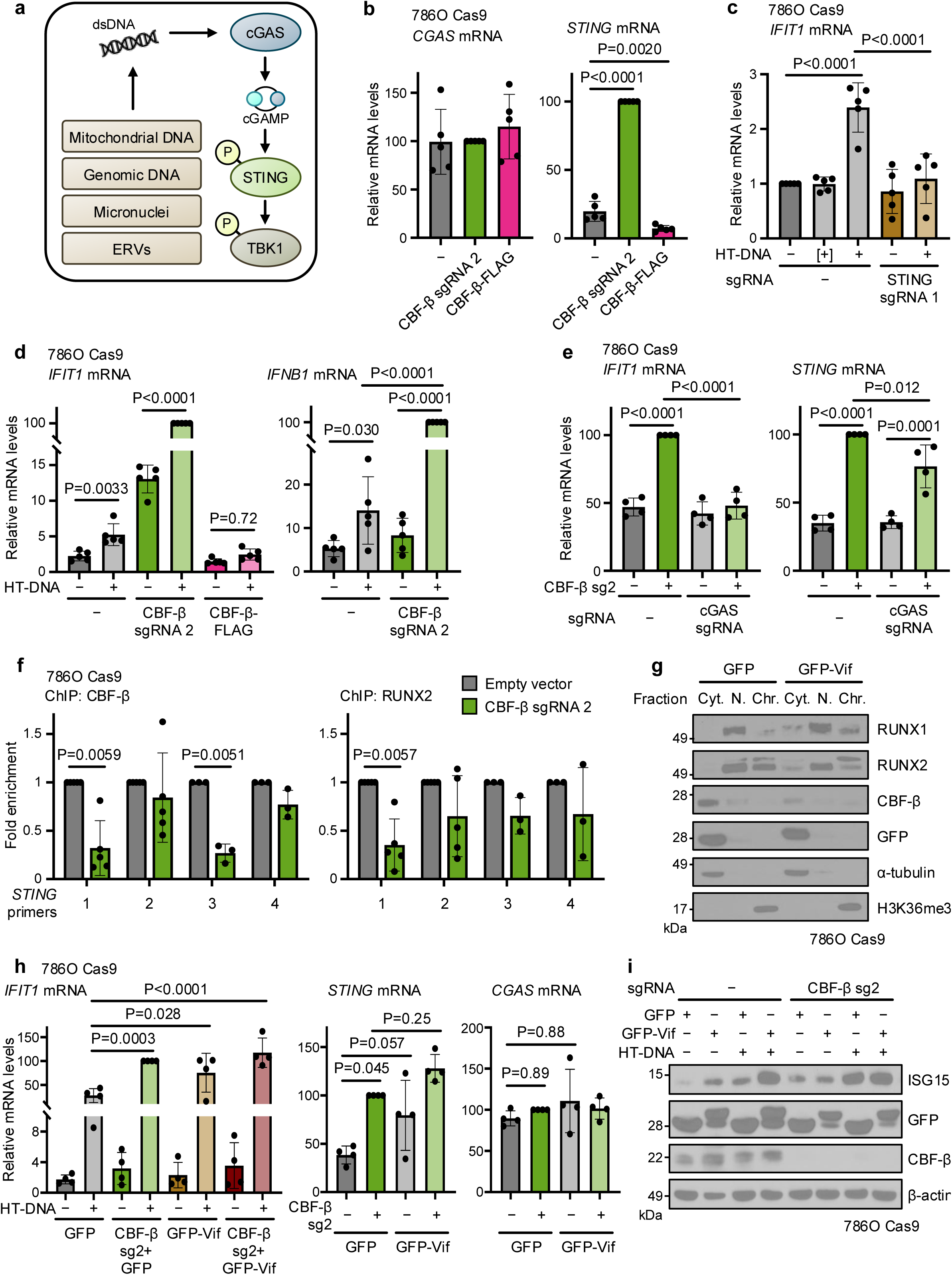
CBF-β represses STING to tune the type I interferon response (a) cGAS-STING signalling axis. cGAS is activated by exogenous or mislocalised double stranded DNA (dsDNA), leading to cGAMP production which triggers a series of phosphorylation events involving STING, TBK1 and IRF3. (b) CBF-β regulates STING transcriptionally. qPCR analysis of *CGAS* and *STING* expression in 786O Cas9 cells transduced with CBF-β sg2, a wild-type CBF-β-FLAG overexpression construct, or an empty lentiviral vector. n=3 biologically independent replicates. Mean ± SD. One-way ANOVA. (c) 786O Cas9 cells were transduced with an sgRNA targeting STING, or an empty vector control, and transfected 6 hours prior to analysis with 0.5 μg/ml herring testis DNA (HT-DNA) as indicated. Additional controls (indicated by [+]) were treated for 6 hours with 0.5 μg/ml HT- DNA, in the absence of the Lipofectamine 2000 transfection reagent. qPCR analysis of *IFIT1* mRNA expression. n=3 biologically independent replicates. Mean ± SD. Two-way ANOVA. (d) CBF-β controls the magnitude of the ISG response to dsDNA transfection. qPCR analysis of 786O Cas9 cells transduced with CBF-β sg2, a wild-type CBF-β-FLAG overexpression construct, or an empty vector, and either transfected with 0.5 μg/ml HT-DNA for 6 hours or left untransfected. n=3 biologically independent replicates. Mean ± SD. Two-way ANOVA. (e) qPCR analysis of 786O Cas9 cells transduced with sgRNAs targeting CBF-β (sg2) and cGAS, or an empty vector. n=4 biologically independent replicates. Mean ± SD. Two-way ANOVA. (f) Chromatin binding of CBF-β and RUNX2 at various sites in the *STING* gene, by ChIP- qPCR analysis of 786O Cas9 cells transduced with CBF-β sg2 or an empty vector. Primer locations are illustrated in **Extended Data** Fig. 6d. n=5 (primers 1 and 2), and n=3 (primers 3 and 4) biologically independent replicates. Mean ± SD. Unpaired *t*-test. (g) Subcellular fractionation following transduction of 786O Cas9 cells with an HIV GFP-Vif expression construct, or a GFP-only control. Cyt.: cytoplasmic fraction. N.: nucleoplasmic fraction. Chr.: chromatin-associated fraction. Immunoblot representative of 2 biologically independent replicates. (**h,i**) 786O Cas9 cells were transduced with vectors encoding CBF-β sg2, GFP-Vif, or a GFP- only control, as indicated, and either transfected with 0.5 μg/ml HT-DNA for 6 hours or left untransfected. qPCR analysis of *IFIT1*, *CGAS* and *STING* mRNA expression (**h**), and immunoblot analysis of ISG15 levels (**i**). n=4 biologically independent replicates. Mean ± SD. Two-way ANOVA.

It was possible that CBF-β loss may contribute to STING activation through the release of dsDNA into the cytoplasm from either a mitochondrial or genomic source^56–62^. However, we found no evidence of mitochondrial dysfunction or superoxide generation by flow cytometry in 786O cells (**Extended Data** Fig. 5a,b), and neither CBF-β deletion nor overexpression affected the cellular oxygen consumption rate (**Extended Data** Fig. 5c). Moreover, there was no overt genomic DNA damage or endogenous retrovirus (ERV) expression in response to transduction with CBF-β sg2 (**Extended Data** Fig. 5d **and Supplementary Table 4**). Indeed, while a basal level of cGAMP was required for STING activity, cGAS knockout did not affect CBF-β knockout-induced STING expression (**Fig. 6e and Extended Data** Fig. 5e). Instead, our results indicated that CBF-β-mediated regulation of STING levels was sufficient to tune a cell-intrinsic ISG response. In support of this, equivalent experiments confirmed the synergistic induction of ISGs and *IFNB1* upon CBF-β depletion and HT-DNA transfection in several other cell lines, including those of non-cancerous renal origin (**Extended Data** Fig. 6a-c).

The involvement of CBF-β in tuning *STING* transcription implied that it may be tonically repressed by CBF-β/RUNX dimers. To interrogate this, we first examined existing chromatin immunoprecipitation (ChIP) sequencing data taken from the chronic myeloid leukaemia K562 cell line, and observed RUNX1 binding at the *STING* promoter region (**Extended Data** Fig. 6d), associated with a 5’-YGYGGTY-3’ RUNX consensus motif at the 5’-end of the gene^63,64^. We generated primers spanning the RUNX binding region at the promoter and putative distal enhancer of STING, and confirmed that CBF-β and RUNX2 associate at the 5’-terminal promoter-like region (primers 1 and 3) (**Fig. 6f and Extended Data** Fig. 6d). Together, these findings indicate that CBF-β tunes the level of STING through transcriptional repression.

Finally, we explored whether the tonic regulation of STING by CBF-β has broader implications for type I interferon responses. CBF-β is implicated in human/simian immunodeficiency virus (HIV/SIV) infection, whereby the virally-encoded Vif associates with CBF-β to hijack a ubiquitin E3 ligase complex and degrade host APOBEC3G^65,66^. Besides disabling this important layer of antiviral defence, the trapping of CBF-β by Vif in the cytoplasm also prevents it from co- operating with RUNX transcription factors and exerting transcriptional activity (**Fig. 6g**)^67^. We therefore expressed GFP-Vif in 786O cells, and measured basal and dsDNA-induced type I interferon signalling. Vif expression increased ISG expression, similarly to CBF-β loss (**Fig. 6h**), and triggered the expression of both STING and downstream ISGs (**Fig. 6h,i**). Together, our findings establish CBF-β as a core and direct regulator of STING-mediated interferon signalling, with broad physiological and pathological implications in the context of anti- retroviral immunity and kidney cancer (**Fig. 7**).

**Fig. 7.**
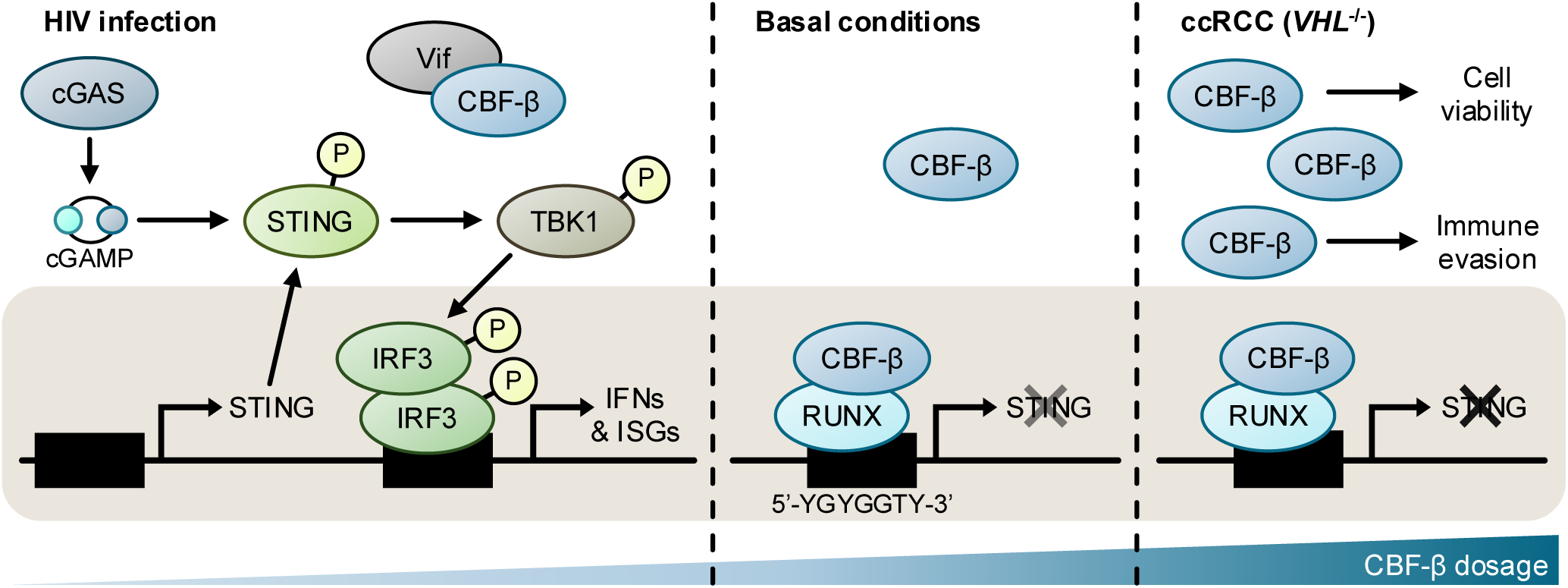
Model of regulation of STING expression by CBF-β/RUNX In basal conditions, CBF-β acts as a rheostat to fine-tune cGAS-STING pathway activity. In the absence of CBF-β/RUNX nuclear translocation, as occurs with sequestration of CBF-β by the lentiviral protein Vif, STING is abundantly expressed, thus amplifying the response to cGAS activation to stimulate signalling through TBK1 and IRF3, ultimately inducing an IFN/ISG-dependent antiviral state. Conversely, CBF-β and RUNX levels are elevated in ccRCC, permitting cell growth. Additionally, CBF-β and RUNX binding at the *STING* locus suppresses IFN signalling and likely promotes immune escape.

## Discussion

The synthetic lethal interaction between *VHL* and *CBFB* provides a new route to target ccRCCs, and is supported by our murine xenograft models in which tumour growth is markedly inhibited. The lethality phenotype following *CBFB* loss is consistent both with observations made upon depletion of RUNX1 and RUNX2 in human kidney cancer lines^68^, and with CBF-β sitting among the top hits of a computational screening platform to identify synthetic lethal interactions of cancers^24^. Prior studies have also observed that high RUNX2 expression in kidney cancer is associated with a mesenchymal phenotype which promotes tumour cell migration, invasion and proliferation *in vitro*, and with higher tumour grade *in vivo*^69–71^. Similarly, elevated RUNX1 in tumour biopsies correlates with poor survival, which has been attributed to altered transcription of extracellular matrix components and remodellers^68,72^. Therefore, although there has been relatively poor agreement between prior VHL-associated synthetic lethality datasets to date^24–31,34^, there is now compelling evidence that CBF-β, along with RUNX proteins, is an emerging therapeutic target for ccRCC.

Whilst both CBF-β and RUNX proteins exert synthetic lethal effects with *VHL* loss, the formation of a heterodimeric complex may not be required, as CBF-β overexpression increased RUNX1/2 levels irrespective of direct binding (**Fig. 3**). This presumed translational effect is consistent with earlier work on CBF-β and RUNX levels in breast cancer cells^73^. The ability of CBF-β to control RUNX levels may also explain prior observations in *Cbfb*-null mouse embryos, in which Runx1 protein was barely detectable despite these embryos displaying similar levels of Runx1 mRNA to *Cbfb*-proficient and -heterozygous controls^74^.

There is also compelling evidence from other systems that RUNX can exert CBF-β- independent roles^75–78^, including the switch between transcriptional activation and repression by RUNX isoforms^41,79–83^. Interestingly, we detected a net increase in global transcription in our RNA sequencing studies upon CBF-β deletion, which was abrogated by *VHL* re- expression. If the primary function of CBF-β in 786O cells is indeed the regulation of RUNX protein abundance, this may indicate that *VHL* loss causes RUNX to adopt a more repressive state. Further dissection of such pathways which modulate RUNX function could therefore unveil new therapeutic targets to specifically modulate cell survival and STING expression in ccRCC while sparing the critical role of RUNX in other lineages.

Kidney cancer cell proliferation relies on the suppression of tumour cell-intrinsic type I IFN activity, and our findings that disruption of CBF-β function can activate a cell-intrinsic ISG response in ccRCC cells provide avenues to overcome this repression. Large deletions within chromosomes 9p and 14q are associated with metastasis and poor prognosis in ccRCC^84^, and the loss of a cluster of type I IFN genes within the affected chromosomal region 9p21.3 has separately been linked to impaired immune surveillance and increased metastasis^85,86^. In addition, mutations in the tumour suppressors *BAP1*, *PBRM1* and *SETD2*, which co-localise with *VHL* on chromosome 3p and are frequently disrupted in ccRCC, suppress ISGF3- mediated ISG expression^87–89^. Therefore, we postulate that targeting CBF-β could enhance tumour regression in response to immune checkpoint inhibition, as has been observed in trials upregulating type I IFN responses through synthetic cyclic dinucleotides and STING agonists^90,91^.

Since the discovery of IRF3 activation by STING, it has been evident that modulation of STING alone, irrespective of cGAS, can be sufficient to stimulate IRF3 phosphorylation^92,93^. Yet despite this, it is striking that only a handful of studies have interrogated the *STING* locus in health and disease to reveal its control by CREB, c-Myc and NF-κB^94–96^. While RUNX1 has been previously shown to attenuate interferon signalling in granulocyte–monocyte progenitors through an unclear mechanism^97^, our studies now implicate RUNX/CBF-β as a rheostat of STING levels. This axis could be exploited not just in cancer but also in the context of a wide range of interferonopathies, including those triggered by pathological sensing of cytosolic ‘self’ DNA or by activating mutations in *STING* itself. Moreover, as STING-driven interferon signalling is an important factor in the clearance of HIV-1^98,99^, the regulation of the STING locus by RUNX/CBF-β may have evolved within mammalian cells as a method to counteract its sequestration by Vif. Understanding how CBF-β and other factors co-operate to tune the balance between appropriate and pathological STING expression remains an important avenue for further research, with broad implications for the innate immune control of tumours and for understanding host responses to viral infections.

## Acknowledgements

We thank all members of the Nathan lab for their helpful comments on the work and manuscript. We also thank Jan Rehwinkel for discussions regarding interferon signalling. This work was supported by a Wellcome Senior Clinical Research Fellowship to JAN (215477/Z/19/Z), a CRUK PhD studentship to JACB, and a Lister Institute Research Fellowship to JAN. The QZ lab was supported by Cancer Prevention and Research Institute of Texas (CPRIT) (RR190058) and VHL synthetic lethality-related research in the QZ lab is partially supported by National Cancer Institute (R01CA284591). AOS was supported by a grant from the Wellcome Sanger Institute and Wellcome Trust (206194) and JJS was supported by a grant from Open Targets (OTAR2080). TJM is supported by a CRUK fellowship (C63474/A27176). This work was also supported by the NIHR BRC.

## Author contributions

Conceptualization, JACB and JAN; Methodology, JACB, TP, JCW, NW, QZ, NJM, BMO, AOS and JAN; Investigation, JACB, TP, QL, NW, JCW, JJS, TJM, AOS, QZ and JAN; Writing – original draft, JACB, TP and JAN; Writing – reviewing and editing, all authors; Funding acquisition, JAN; Resources, NJM, AOS, QZ and JAN; Supervision, BMO, AOS, QZ and JAN.

## Methods

### Cell lines

Cell lines and reagents were sourced as detailed in **Supplementary Table 5**. 786O, 786O+*VHL* and 769P cells were cultured in Roswell Park Memorial Institute 1640 medium (RPMI-1640), and A498, HEK293T, HKC8, HK2, RCC4, RCC4+*VHL* and RCC10 cells were cultured in Dulbecco’s Modified Eagle Medium (DMEM). All media was supplemented with 10% Fetal Calf Serum (FCS). Cells were maintained at 37°C in 5% CO_2_. 2 μg/ml or 2.5 μg/ml Puromycin, 5 μg/ml Blasticidin, or 200 μg/ml Hygromycin was added to the media for antibiotic selection of transduced and transfected cells. Hypoxic culture was undertaken in a Whitley H35 Hypoxystation (Don Whitley) set at 1% O_2_, 5% CO_2_ and 94% N_2_. Cells were confirmed as mycoplasma-free with the MycoAlert detection kit (Lonza Cat#LT07-318), and authenticated by short tandem repeat profiling (Eurofins Genomics).

### Plasmids

Cloning of novel gene expression plasmids was performed using Gibson assembly (NEB) of PCR amplicons and restriction digested backbone vectors, using the DNA sequences indicated in **Supplementary Table 6**. Plasmids were amplified in NEB 5α Competent *E. coli* (Cat#C2987H) grown at 37°C in LB containing appropriate antibiotics. The sources of plasmids used in this study are indicated in **Supplementary Table 5**.

### Lentiviral production and transduction

Lentivirus was produced by transfection of HEK293T cells with the relevant vector, pMD.G, and pCMVR8.91 in six-well plates or 15 cm dishes at 60-70% confluence using Fugene 6 Transfection Reagent and Opti-MEM I. Viral supernatant was collected after 48 hours through a 0.45 μm filter and stored at −80°C. For small-scale transductions, 200 μl virus was applied to 5×10^4^ cells in 24-well plates containing 500 μl media, and appropriate antibiotic selection performed after 48 hours until all untransduced control cells had died.

Lentiviral expression constructs were used for gene overexpression. Assays were performed 10-14 days after transduction in comparison to controls transduced with the pHRSIN-SFFV- Puro empty vector.

### CRISPR-Cas9 targeted gene deletions

Gene-specific sgRNA sequences listed in **Supplementary Table 6** were designed using the E-CRISP or Vienna Biocenter algorithms^101,102^. For cloning into the pKLV-U6gRNA(BbsI)- PGKpuro2ABFP, pKLV-U6gRNA(BbsI)-PGKblast2ABFP and pSpCas9(BB)-T2A-Puro vectors, sgRNA oligonucleotides were designed with appropriate overhangs and cloned according to the Zhang lab protocol^103^. Cloning into tet-pLKO-sgRNA-puro for doxycycline-inducible sgRNA expression was performed using modified overhangs using the method of Huang et al.^104^. Multiplexed expression of sgRNAs targeting RUNX1, RUNX2 and RUNX3, or STAT1 and STAT2 from the same pKLV-U6gRNA(BbsI)-PGKpuro2ABFP vector was achieved by PCR amplification of the tRNA scaffold of pCFD5 with primers containing the relevant sgRNA sequences, followed by Gibson assembly^105^.

Cells were transduced with Lenti-Cas9-2A-Blast, or had previously been transduced with an equivalent Hygromycin-resistant vector^106^, to stably express Cas9. To generate stable CRISPR/Cas9 deletion mutants, sgRNAs expressed from pSpCas9(BB)-T2A-Puro were transiently transfected into cells as described above, before dilution cloning in flat-bottomed 96-well plates. For other CRISPR/Cas9 experiments, cells were transduced with the indicated sgRNA expressed from pKLV-U6gRNA(BbsI)-PGKpuro2ABFP, pKLV-U6gRNA(BbsI)-

PGKblast2ABFP, or tet-pLKO-sgRNA-puro, and assays performed 10-14 days after transduction, following antibiotic selection. All CRISPR/Cas9 experiments were controlled by transduction with an equivalent empty vector.

### Pooled genome-wide CRISPR/Cas9 screening

At least 1.2×10^8^ Cas9-expressing cells of 786O, 786O+*VHL*, RCC4 and RCC4+*VHL* backgrounds were transduced with the TKOv3 library at a multiplicity of infection of approximately 30%. Cells were expanded to 15 cm dishes after 27 hours, and selected with Puromycin for seven days. Every two or three days, cells were pooled, counted and passaged. After 17 population doublings, genomic DNA was extracted from 4×10^7^ cells using a Puregene Cell Kit (Qiagen Cat#158043). Lentiviral sgRNA inserts were amplified in a two-step PCR using the primers detailed in **Supplementary Table 6**, with the second step introducing unique barcode sequences for multiplexed sequencing^107^. Amplicons were cleaned with Agencourt AMPure XP beads and sequenced on HiSeq 4000 or NovaSeq 6000 (Illumina) with a custom sequencing primer (**Supplementary Table 6**). An average of at least 400-fold (786O screen) or 500-fold (RCC4 screen) representation of each sgRNA within the pool was maintained throughout all stages of the screens.

Sequencing data was processed and analysed using Cutadapt, HISAT2 and Bayesian Analysis of Gene Essentiality version 2 (BAGEL2) in a Python-based bioinformatic pipeline. Read counts for each sgRNA were compared across paired *VHL*-proficient and -deficient cell lines, and between late and early timepoints in each cell line, computing Bayes Factors and False Discovery Rates (FDRs).

### shRNA-mediated RNA depletion

Oligonucleotides were designed with a TTCAAGAGA hairpin, using shRNA sequences from the Broad Institute RNAi Consortium shRNA Library (**Supplementary Table 6**). Sequences were cloned into pC.SIREN.Puro by digestion and ligation. Cells were transduced with the indicated vector or a scrambled shRNA control, and assays performed after at least seven days, following Puromycin selection.

### Proliferation and cell death assays

Cells were seeded in six-well plates at predefined densities (1.5×10^4^ cells/well for 786O cells, 2.5×10^4^ cells/well for other cell lines), and manually counted with a haemocytometer after 96 hours. For the IFN-β dosage experiment, cells were seeded with the indicated concentration of IFN-β at 1×10^4^ cells/well in 12-well plates, and final counts after 72 hours were normalised to an untreated control.

### Clonogenic assay

Cells were seeded in six-well plates at a density of 100 cells/well and incubated undisturbed for seven days. Colonies were fixed with 100% ice-cold methanol for 10 minutes, visualised with 0.5% crystal violet in 25% methanol, and counted.

### Flow cytometry

Following appropriate staining, cells were collected in 5 ml FACS tubes, centrifuged, washed and resuspended in PBS prior to analysis on a Fortessa flow cytometer (BD Biosciences) using FlowJo software. Gating strategies are displayed in **Supplementary** Fig. 2.

Cell death experiments employing SYTOX AADvanced and CellEvent Caspase-3/7 Green were performed according to the manufacturer’s instructions, with gates set based on empty vector-transduced or DMSO treated controls. For cell death inhibitor treatments, cells were seeded at a density of 2×10^4^ cells/well in 12-well plates in media containing the indicated drug concentration or a DMSO vehicle. The media was exchanged after 24 hours with fresh media containing the same dose of inhibitor, and analysis performed as above after a further 24 hours.

To assess mitochondrial function, cells were seeded in six-well plates at 8×10^4^ cells/well 24 hours prior to analysis with MitoSOX Red, MitoTracker Green FM, and tetramethylrhodamine methyl ester (TMRM). Before staining, control wells were treated with 30 minutes of Carbonyl cyanide-p-trifluoromethoxyphenylhydrazone (FCCP) or Antimycin A. For MitoTracker Green and TMRM staining, cells were harvested, centrifuged and stained in FACS tubes for 30 minutes at 37°C. MitoSOX Red staining was performed at 37°C for 10 minutes directly in six-well plates, having washed cells in Hanks’ Balanced Salt Solution (HBSS).

### Competitive growth assay

Seven days post-transduction and following antibiotic selection, 3×10^4^ clonal Cas9-expressing 786O mCherry cells were mixed with an equal number of clonal Cas9-expressing 786O+*VHL* GFP cells in six-well plates. For doxycycline-inducible sgRNA expression vectors, 100 ng/ml doxycycline was applied for nine days following the establishment of the co-culture. At each passage, a subset of each culture was harvested in 3.6% paraformaldehyde for flow cytometry analysis. The 786O mCherry:786O+*VHL* GFP cell ratio was normalised to 0.5 for the first time- point, and then to that of a relevant empty vector or scrambled shRNA control.

### Cell cycle analysis

BrdU incorporation and propidium iodide staining of total cellular DNA content were used to assay cell cycle phases. For dual staining, 3.5×10^5^ cells were seeded in 10 cm dishes. After 24 hours, cells were incubated in media containing 10 μM BrdU for one hour.

For intracellular staining of γ-H2A.X, cells were permeabilised with ice-cold 90% methanol for 30 minutes, treated with 2 M HCl for 30 minutes at room temperature, incubated at 4°C for 30 minutes with the indicated dilution of primary antibody in a 1% BSA/PBS incubation buffer, washed in PBS, and stained with a secondary antibody at 4°C for a further 30 minutes. Cells were finally incubated in 3 μM propidium iodide/PBS for 15 minutes before flow cytometry analysis.

Cell cycle synchronisation was achieved using a double thymidine block method. 3.5×10^4^ cells were seeded in six-well plates. Cells were treated with 2 μM thymidine after seven hours, which was replaced with thymidine-free media after 15 hours. After eight hours, media was replaced again with 2 μM thymidine for 17 hours. At the end of this second treatment, and for every two hours thereafter for 14 hours, samples were harvested in 90% methanol before propidium iodide staining and flow cytometry analysis. An asynchronous control of each experimental condition was maintained in thymidine-free media throughout.

### LDH release cytotoxicity assay

5×10^3^ cells were seeded in triplicate in 96-well plates with 100 μl of media. After 24 hours, 50 μl supernatant from each well was transferred to a new plate, and the Pierce LDH Cytotoxicity Assay Kit (ThermoFisher Cat#88953) used to assess LDH levels as per the manufacturer’s instructions, with absorbance measurements taken using a CLARIOstar Plus plate reader (BMG Labtech).

### DNA transfection

8×10^4^ 786O Cas9 cells were seeded in 2 ml media in six-well plates 24 hours prior to transfection with herring testes DNA (HT-DNA). Mixes of 4 μg/ml HT-DNA and 40 μg/ml Lipofectamine 2000, prepared in Opti-MEM I, were combined and gently mixed. After 30 minutes incubation at room temperature, 500 μl DNA/Lipofectamine 2000 complex was added dropwise to cells.

### Real-time quantitative PCR (qPCR)

Cell lysis was performed in six-well plates at approximately 70-80% confluence. Total RNA was extracted and purified with the RNeasy Plus Mini Kit (Qiagen Cat#74136), and cDNA generated using Protoscript II Reverse Transcriptase. 20 ng template cDNA was amplified with SYBR Green PCR Master Mix on a QuantStudio 7 Real-Time PCR System (ThermoFisher), using primers specific to the mRNA interest (**Supplementary Table 6**). qPCR reactions for each cDNA/primer combination were performed in technical triplicates, and the mean used to calculate the relative transcript abundance with the ΔΔCt method following normalisation to the reference index of a housekeeping gene (*β-Actin*).

### RNA sequencing

Total RNA was extracted using the PureLink RNA Mini Kit (ThermoFisher Cat#12183025), including on-column DNA digestion with PureLink DNase. Library preparation and sequencing were undertaken by GENEWIZ (Azenta), using NEBNext Ultra II RNA Library Preparation Kit (NEB Cat#E7770L) for polyA transcript selection, and NovaSeq 6000 (Illumina). Bioinformatic analysis of RNA-Seq data with DESeq2 was performed as described previously^106^, except using Salmon for read mapping and STAR/TEtranscripts for differential expression analysis of transposable elements. Gene Set Enrichment Analysis of Hallmark gene sets was undertaken in R using the fgsea package.

### ChIP-qPCR

786O cells grown to 70% confluence in 15 cm dishes were treated with 1% formaldehyde for 10 minutes to crosslink proteins to chromatin, and the reaction quenched with 0.125 M glycine for 10 minutes at room temperature. Cells were washed twice in ice-cold PBS, transferred to tubes, centrifuged at 800 rpm for 10 minutes and lysed in 500 µl of ChIP lysis buffer (50 mM Tris-HCl pH 8.1, 1% SDS, 10 mM EDTA, and 1×cOmplete EDTA-free protease inhibitor cocktail). Next, samples were incubated for 10 minutes on ice, and diluted with an equivalent volume of ChIP dilution buffer (20 mM Tris-HCl pH 8.1, 1% (v/v) Triton X-100, 2 mM EDTA, and 150 mM NaCl) prior to sonication using beads in a Bioruptor (Diagenode) for 20 cycles of 15 seconds on and 30 seconds off. Samples were centrifuged at 4°C for 10 minutes at 13,000 rpm and supernatants collected and stored at −20°C as input controls. 200 µl aliquots of the remaining samples were diluted with ChIP dilution buffer to 1 ml, and precleared with 25 µl Protein G magnetic beads at 4°C for two hours before overnight immunoprecipitation with the indicated primary antibody at 4°C. Next, 25 μl Protein G magnetic beads were added and samples incubated for a further two hours at 4°C. Beads were washed for 5 minutes with wash buffer 1 (20 mM Tris-HCl pH 8.1, 0.1% (w/v) SDS, 1% (v/v) Triton X-100, 2 mM EDTA, and 150 mM NaCl), wash buffer 2 (as for wash buffer 1 with 500 mM NaCl), wash buffer 3 (10 mM Tris-HCl pH 8.1, 0.25 M LiCl, 7 1% (v/v) NP-40, 1% (w/v) Na deoxycholate, and 1 mM EDTA), and twice with TE buffer (10 mM Tris-HCl pH 8.0, and 1 mM EDTA). Complexes were eluted in 120 μl elution buffer (1% (w/v) SDS and 0.1 M NaHCO_3_), and incubated overnight at 65°C with agitation with 0.2 M NaCl to reverse the formaldehyde crosslinks. Samples were incubated with 20 μg Proteinase K for 4 hours at 45°C, and with RNase H for 30 minutes at 37°C to digest protein and RNA, before DNA was purified using the DNA MinElute kit (Qiagen Cat#28206). DNA was analysed by qPCR, and results were normalised to the amplification of the input material for each sample.

### Analysis of ChIP sequencing data

The *STING* locus was visualised on the RefSeq hg38 (GRCh38) human genome build using the Integrative Genomics Viewer (IGV), with annotated transcripts derived from the Ensembl genome browser release 94^108^. RUNX1 ChIP-Seq data (https://www.encodeproject.org/files/ENCFF985UVT) and candidate *cis*-regulatory element annotations (https://www.encodeproject.org/files/ENCFF657KYN) were extracted from ENCODE data files, using the ENCODE data portal.

### Liquid-chromatography mass spectrometry (LC-MS)

2×10^6^ cells were harvested in PBS, resuspended in 71 μl resuspension buffer (76 mM triethylammonium bicarbonate (TEAB) pH 8.5, 3 mM MgCl2, 1,400 U/ml benzonase, 7 mM tris(2-carboxyethyl)phosphine (TCEP), and 28 mM chloroacetamide), and lysed with the addition of 25 μl 20% SDS for 30 minutes at 25°C in the dark. 4 μl 1.25 M Azido-PEG3-Azide was added and samples incubated for 20 minutes at 55°C to quench the TCEP. 5 μl aliquots were compared to a standard curve of BSA using a Pierce Microplate BSA assay (ThermoFisher Cat#23252), and samples were normalised to 25 μg by dilution in a 50 mM TEAB, 5% SDS buffer.

To complete the denaturation, a 10% volume of 27.5% phosphoric acid was added to each sample, causing acidification to approximately pH 2. A 6× volume of a wash buffer (100 mM HEPES pH 7.55, 90% methanol) was added, and the solution transferred to an S-trap micro column using a positive pressure manifold (Tecan M10) and in-house fabricated adaptors. Samples were washed with 1:1 (v/v) methanol:chloroform and three times with the wash buffer. S-traps were centrifuged at 4,000 g for 2 minutes to remove residual wash buffer, and 30 μl of a digestion solution (50 mM TEAB pH 8.5, 0.1% Na deoxycholate) containing 1.25 μg Trypsin/lysC Mix was added. S-traps were incubated for 16 hours at 37°C, and peptides recovered by the addition of 25 μl digestion solution and incubation at room temperature for 15 minutes, before centrifugation at 4000 g. 40 μl 0.2% formic acid (FA) and 25 μl 50% acetonitrile (ACN) were used in the same manner to elute samples, which were subsequently dried in a vacuum centrifuge equipped with a cold trap. Samples were resuspended in 21 μl 100 mM TEAB pH 8.5. 0.5 μg unique TMTpro labels, resuspended in 9 μl anhydrous ACN, were added to each sample and incubated at room temperature for one hour. To confirm labelling efficiency of >98% and equal loading, 3 μl aliquots of each sample were taken, pooled, and analysed by LC-MS.

Samples were normalised by adjusting for the total reporter ion intensities of the initial test, and were pooled and dried in a vacuum centrifuge. Samples were acidified with approximately 200 μl 0.1% Trifluoroacetic Acid (TFA), and FA was added until the Na deoxycholate was visibly precipitated. Four volumes of ethyl acetate were added and vortexed vigorously for 10 seconds. Samples were centrifuged at 15,000 g for 5 minutes at room temperature for phase separation, and the aqueous phase transferred to a fresh tube. Samples were partially dried in a vacuum centrifuge and made up to 1 ml with 0.1% TFA. FA was added to achieve a pH <2, and the samples cleaned by solid phase extraction using a 50 mg tC18 Sep-Pak cartridge (Waters) and a positive pressure manifold: the cartridge was wetted with 1 ml methanol followed by 1 ml ACN, equilibrated with 1 ml 0.1% TFA, and each sample loaded slowly. The cartridge was washed with 1 ml 0.1% TFA, and samples were eluted in 750 μl 80% ACN, 0.1% TFA and dried under a vacuum.

Next, samples were resuspended in 40 μl ammonium formate pH 10 and transferred to a glass HPLC vial. Basic pH reverse-phase fractionation was conducted on an Ultimate 3000 UHPLC system (ThermoFisher) equipped with a 2.1 mm × 15 cm, 1.7 μm Kinetex EVO column (Phenomenex) set at a flow rate of 500 μl/minute, using solvent A (3% ACN), solvent B (100% ACN), and solvent C (200 mM ammonium formate pH 10; kept at a constant 10%). Samples were loaded in 90% solvent A for 10 minutes before they were eluted in a gradient of solvent B of 0-10% over 10 minutes, 10-34% over 21 minutes, and 34-50% over 5 minutes, followed by a 10 minute wash with 90% solvent B. 100 μl fractions were collected throughout the run, and those containing peptide (determined by ultraviolet absorbance at 280 nm) were recombined across the gradient to preserve orthogonality with on-line low pH reverse-phase separation. Combined samples were dried in a vacuum centrifuge and stored at −20°C.

Samples were analysed by LC-MS on an Orbitrap Fusion instrument on-line with an Ultimate 3000 RSLC nano UHPLC system (ThermoFisher), using trapping solvent (0.1% TFA), analytical solvent A (0.1% FA) and analytical solvent B (0.1% FA in ACN). After resuspension in 10 μl 1% TFA and 5% DMSO, 5 μl of each sample was loaded onto a 300 μm × 5 mm PepMap cartridge trap (ThermoFisher) at 10 μl/minute for 5 minutes. Samples were separated on a 75 cm × 75 μm I.D. 2 μm particle size PepMap C18 column (ThermoFisher) with a gradient of analytical solvent B at 3-10% over 10 minutes, 10-35% over 155 minutes, and 35- 45% over 9 minutes, followed by washing with 95% analytical solvent B for 5 minutes and re-equilibration at 3% analytical solvent B. Eluted particles were next introduced to the MS by 2.1 kV electrospray to a 5 cm × 30 μm stainless steel emitter (PepSep). The MS was operated in SPS mode, in which MS1 Scans are obtained in the Orbitrap, CID-MS2 Scans in the Ion Trap and HCD-MS3 acquired in the Orbitrap to resolve reporter ions.

Data were analysed with Peaks 11, and .raw files searched against the SwissProt Human Database with appended common contaminants. The peptide-spectrum match FDR was controlled at 0.1% by decoy database search. Statistical analysis of the relative abundance of identified proteins in each sample was performed with the limma package in R by a moderated *t*-test. A q-value was computed to determine appropriate cut-off values by correcting P-values for multiple hypothesis testing with the Benjamini-Hochberg method.

### SDS-PAGE immunoblotting

Cells were lysed at approximately 70-80% confluence in six-well plates with an SDS loading buffer (1% SDS, 50 mM Tris pH 7.4, 150 mM NaCl, 10% glycerol, and 2 μl/ml benzonase) and incubated on ice for 20 minutes before heating at 90°C for 5 minutes. Proteins were resolved by SDS-PAGE (Bio-Rad), and transferred to methanol-activated polyvinylidenedifluoride (PVDF) membranes. Membranes were blocked with 0.2% TWEEN-20/PBS containing 5% skimmed milk powder and 1% BSA, probed with the appropriate primary and secondary antibodies for at least one hour each, and developed with Pierce Enhanced Chemiluminescent, SuperSignal West Pico Plus Chemiluminescent, or SuperSignal West Dura Extended Duration substrates. PBS was substituted with TBS for the detection of phosphorylated proteins. β-Actin levels were used to confirm the equivalent loading of each sample. Indicated protein sizes were determined by comparison to a SeeBlue Plus2 pre- stained protein standard.

### Immunoprecipitation

786O cells were lysed in 1 ml 1% Triton X-100/TBS supplemented with 1×cOmplete EDTA-free protease inhibitor cocktail at 4°C for 30 minutes. Lysates were centrifuged at 14,000 g for 15 minutes at 4°C, and the supernatant collected. 20 μl of each sample was taken in an equal volume 2×SDS loading buffer as an input, and the remainder pre-cleared with Pierce Protein G magnetic beads for one hour at 4°C. Supernatants were incubated overnight with 2 μl of the indicated primary antibody at 4°C with rotation. Next, Pierce Protein G magnetic beads were added for two hours, and samples were washed three times in 1% Triton X-100/TBS lysis buffer. Bound proteins were eluted in 2×SDS loading buffer and analysed by SDS-PAGE immunoblotting.

### Subcellular fractionation

2×10^6^ cells were lysed and incubated with rotation in buffer A (0.1% IGEPAL CA-630, 10 mM HEPES, 1.5 mM MgCl_2_, 10 mM KCl, 0.5 mM DTT, and 1×cOmplete EDTA-free protease inhibitor cocktail) for 10 minutes at 4°C before centrifugation at 1,400 g for 4 minutes at 4°C. The supernatant, representing the cytoplasmic fraction, was collected in 2×SDS loading buffer. The nuclear pellet was washed in buffer A, with centrifugation as before, and resuspended in buffer B (20 mM HEPES, 1.5 mM MgCl_2_, 300 mM NaCl, 0.5 mM DTT, 25% glycerol, 0.2 mM EDTA, and 1×cOmplete EDTA-free protease inhibitor cocktail) for 10 minutes at 4°C. Samples were centrifuged at 1,700 g for 4 minutes at 4°C, and the soluble nucleoplasmic fraction and insoluble chromatin pellet were each collected and resuspended in 2×SDS loading buffer. Subcellular fractions were analysed by SDS-PAGE immunoblotting.

### Bioenergetic analysis

Oxygen consumption rates (OCRs) were measured using a Seahorse XF analyser in order to calculate basal respiration. We performed the Mito Stress Test according to the manufacturer’s instructions using Seahorse XF FluxPak consumables (Agilent Technologies). 1.2 × 10^4^ cells were plated in FluxPak 96-well plates and incubated at 37°C for 24 hours. The medium was replaced with Seahorse XF RPMI, pH 7.4 supplemented with 1 mM sodium pyruvate, 2 mM L-glutamine, and 10 mM glucose, and cells were maintained at 37°C in a non-CO_2_ incubator for 1 hour before the run. Analysis was performed using the program settings: mix for 3 minutes and measure for 3 minutes ×3; inject 1.5 μM oligomycin; mix for 3 minutes and measure for 3 minutes ×3; inject 1 μM FCCP; mix for 3 minutes and measure for 3 minutes ×3; inject 1 µM Rotenone, 1 µM Antimycin A and 2.5 μM Hoechst 33342; mix for 3 minutes and measure for 3 minutes ×3. Basal respiration was calculated by subtracting the OCR following Rotenone and Antimycin A treatment from the basal OCR. Following the assay, results were normalised for cell number by quantifying Hoescht 33342 staining using a CLARIOstar Plate Reader at 355-20/455-30 nm.

### Analysis of TCGA expression and survival data

*CBFB* mRNA expression and survival data for ccRCC tumours was obtained from The Cancer Genome Atlas (TCGA)^7^. The R package survminer was used to perform a log-rank test and plot a Kaplan-Meier curve.

### Single cell expression of interferon pathway genes

Single cell data was downloaded from Li et al. (https://data.mendeley.com/datasets/g67bkbnhhg/1)^46^. The anndata file was first converted to a Seurat object using the sceasy package in R_4.4.0 (https://github.com/cellgeni/sceasy). The single cell expression data was normalised and variance stabilised using regularised negative binomial regression using the SCTransform function from the Seurat_5.1.0 R package. Cell types annotated as either ‘Low quality’ or ‘Unknown’ were removed with the remaining cells collated into their broad principal lineages of epithelial cells, RCC cells, NK cells, myeloid cells, fibroblasts, endothelial cells, T cells and B cells. The average normalised expression data for these cell types in relation to interferon and STING pathway genes were plotted using the DotPlot function in R.

### Tumour xenografts

Animal experiments were performed according to protocols approved by either the University of Cambridge Animal Welfare and Ethical Review Board in compliance with the Animals (Scientific Procedures) Act 1986 and UK Home Office regulations, or by the Institutional Animal Care and Use Committee of UT Southwestern Medical Center following NIH guidelines.

Female NSG mice (NOD.Cg-*Prkdc^scid^ Il2rg^tm1Wjl^*/SzJ (Charles River RRID:IMSR_JAX:005557)) used for subcutaneous xenograft were housed at a density of five animals per individually ventilated cage with *ad libitum* access to food and water in a specific pathogen free unit. 786O Cas9 cells expressing the indicated sgRNAs were harvested and washed in Dulbecco’s PBS without calcium and magnesium (D-PBS), and 1×10^7^ cells were administered via subcutaneous injection to the dorsal flank of NSG mice in a blinded and randomised manner between cages. After 28 days, tumour area was determined using callipers as the product of the maximum length and breadth, and mice were euthanised.

Orthotopic xenografts of 786O Cas9 cells stably expressing luciferase and doxycycline-inducible sgRNA vectors were performed in 6-8 week old male and female NSG mice (UTSW Animal Resource Center) as described previously^44^. 1×10^6^ viable cells were resuspended in 20 μL PBS containing 50% Matrigel and injected orthotopically into the left kidney of each NSG mouse. Following successful implantation, mice were fed Purina rodent chow #5001 with 2000 ppm doxycycline to induce sgRNA expression. Tumour growth was monitored weekly by bioluminescence imaging using the SPECTRAL AMI-HTX system. Mice were euthanised after a further eight weeks, and the tumour mass calculated by subtracting the weight of each right kidney from the corresponding left kidney. Metastasis was evaluated in isolated lungs by bioluminescence.

### Quantification and statistical analysis

Quantitative data are expressed as the mean of biological repeats ± 1 standard deviation (SD) or ± 1 standard error of the mean (SEM). P-values were calculated using two-tailed Student’s *t*-tests, two-tailed Mann-Whitney U tests, or analysis of variance (ANOVA), as indicated in figure legends. Statistical analyses of CRISPR/Cas9 screens, RNA-Seq, mass spectrometry, and TCGA data are described in the relevant method sections. The number of biologically independent repeats (independent transductions for experiments involving cell culture, or number of mice for xenograft experiments) are specified in figure legends. Data presented from CRISPR/Cas9 screens are derived from a single replicate in each cell line. Figures were prepared and statistical analyses performed using GraphPad Prism or R. Software used in this study is indicated in **Supplementary Table 5**.

## Data availability

Raw data from RNA sequencing and CRISPR/Cas9 screens have been deposited at GEO and are publicly available at GSE270775 (access token: anwvmykobbsttcl) and GSE270776 (access token: upszseuqvbyddsp). Additional data reported in the paper is available from the lead contact upon request.

Any additional information required to reanalyse the data reported in this paper is available from the corresponding author upon request.

## Code availability

Code for the CRISPR screen analysis pipeline is available at https://github.com/niekwit/crispr-screens (10.5281/zenodo.10286661), the differential transcript analysis pipeline with Salmon/DESeq2 at https://github.com/niekwit/rna-seq-salmon-deseq2 (10.5281/zenodo.10139567), and TE analysis pipeline at https://github.com/niekwit/rna-seq-star-tetranscripts  (10.5281/zenodo.10027278).

---

**Extended Data Fig. 1.**
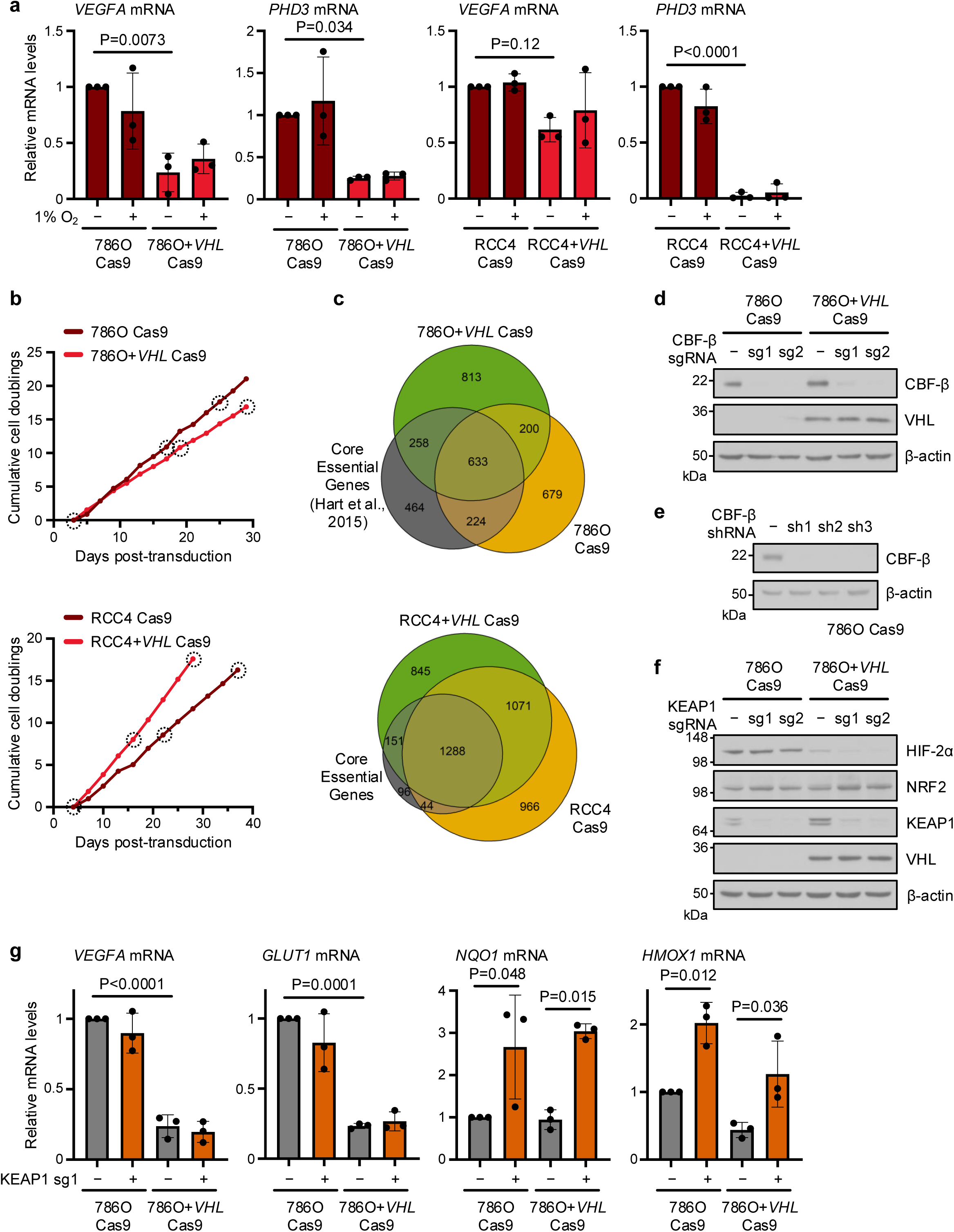
CRISPR/Cas9 screening in ccRCC cell lines reveals *VHL*-associated synthetic lethality of the transcription factors CBF-β and NRF2 (a) Expression of HIF target genes in ccRCC cells and paired VHL-reconstituted cells after culture at 1% or 21% O_2_ for 24 hours. n=3 biologically independent replicates. Mean ± SD. Two-way ANOVA. (b) Cumulative cell doublings from day 3 (786O screen) or day 4 (RCC4 screen) in CRISPR/Cas9 screens. Dotted circles indicate the analysed samples. (c) Efficient identification of essential genes in CRISPR/Cas9 screens. Venn diagrams of essential genes which dropout between early and late timepoints identified by BAGEL2 with FDR<0.05, compared to a reference set of Core Essential Genes^38^. (**d**-**f**) Immunoblots of 786O Cas9 and 786O+*VHL* Cas9 cells transduced with sgRNAs targeting CBF-β (**d**), shRNAs targeting CBF-β (**e**), and sgRNAs targeting KEAP1 (**f**). sgRNA vectors were doxycycline-inducible, and cells were treated with 100 ng/ml doxycycline prior to analysis. Controls were transduced with an empty sgRNA expression vector (**d,f**), or a scrambled shRNA sequence (**e**). Immunoblots representative of 3 independent experiments. (**g**) *KEAP1* knockout induces NRF2, but not HIF, activation. qPCR analysis of HIF targets (*VEGFA* and *GLUT1*) and NRF2 targets (*NQO1* and *HMOX1*) in cells transduced with an sgRNA targeting KEAP1. n=3 biologically independent replicates. Mean ± SD. Two-way ANOVA.

**Extended Data Fig. 2.**
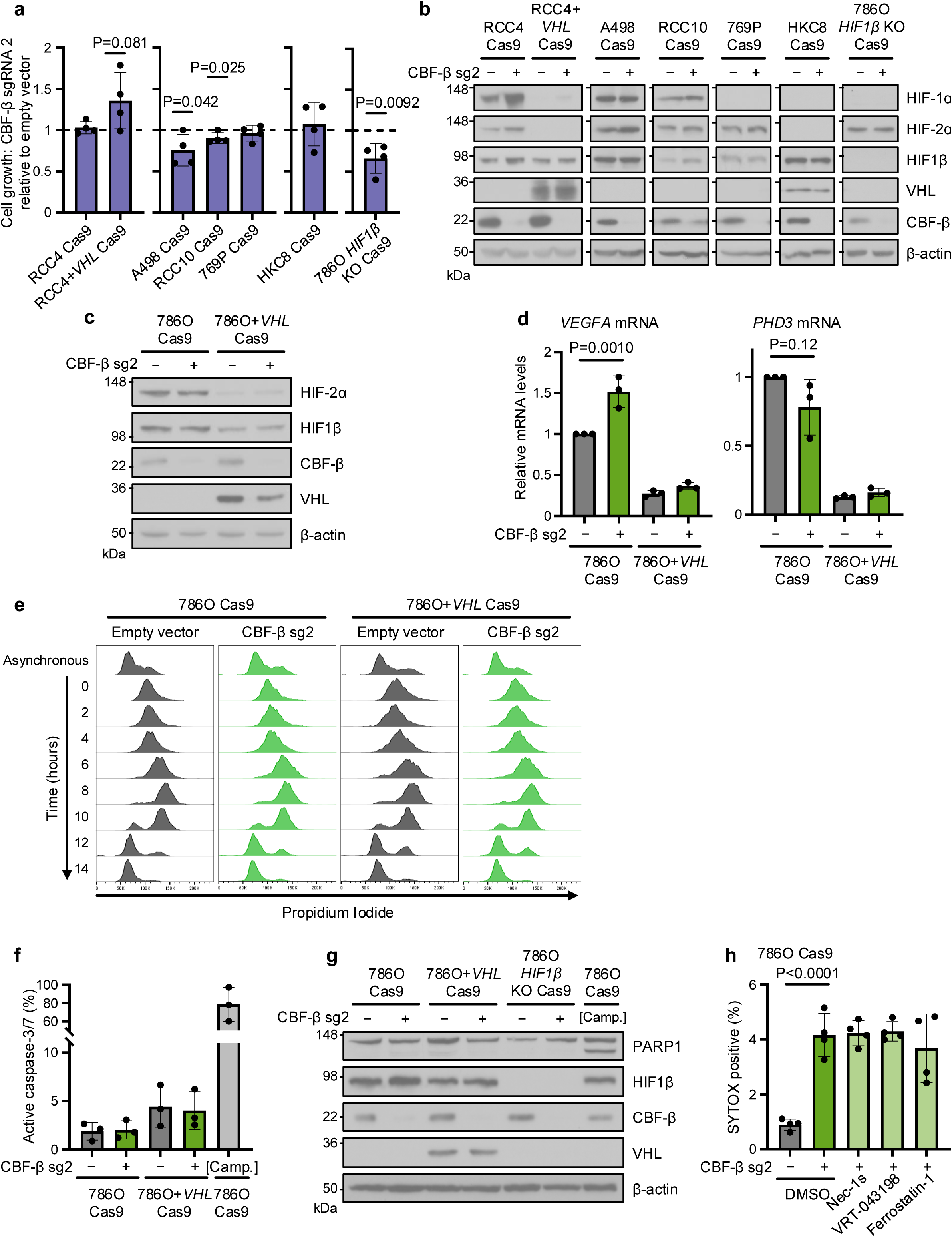
*CBFB* knockout induces cell death in *VHL*-deficient cells (a) Proliferation assay of ccRCC cells (RCC4, A498, RCC10 and 769P), renal proximal tubule epithelial cells (HKC8), and clonal *HIF1β*-deficient 786O cells and *VHL*-reconstituted RCC4 cells, transduced with CBF-β sg2. Cell numbers at day 4 plotted relative to control cultures transduced with an empty vector. n=4 biologically independent replicates. Mean ± SD. Unpaired *t*-test. (b) Representative immunoblot of transductions assayed in **Extended Data** Fig. 2a. n=3 biologically independent replicates. (**c,d**) *CBFB* does not regulate HIF activity in 786O Cas9 cells. 786O Cas9 and 786O+*VHL* Cas9 populations were transduced with CBF-β sg2 or an empty vector control and assayed by immunoblot (**c**), and by qPCR of HIF target genes (**d**). n=3 biologically independent replicates for **c** and **d**. Mean ± SD. Two-way ANOVA. (e) Rate of cell cycle progression in 786O Cas9 and 786O+*VHL* Cas9 cells transduced with CBF-β sg2 or an empty vector control. Cells were synchronised with a double thymidine block, and fixed for propidium iodide staining and flow cytometry every two hours following release. An asynchronous control is provided for comparison. Representative of 3 biologically independent replicates. (f) Proportion of 786O Cas9 and 786O+*VHL* Cas9 cells exhibiting caspase-3/7 activity following transduction with CBF-β sg2 or an empty vector control, as detected by the CellEvent Caspase-3/7 Green flow cytometry reagent. [Camp.]: cells treated with 24 hours 100 μM Camptothecin as a positive control for apoptosis. n=3 biologically independent replicates. Mean ± SD. Two-way ANOVA. (g) PARP1 cleavage in cells transduced with CBF-β sg2 or an empty vector control. Lower band on PARP1 membrane represents the cleaved form of PARP1. [Camp.]: 786O Cas9 cells treated with 10 μM Camptothecin as a positive control for apoptosis. Immunoblot representative of 3 biologically independent replicates. 786O Cas9 cells transduced with CBF-β sg2 or an empty vector control, and treated for 48 hours with 50 μM Necrostatin-1s (Nec-1s), 25 μM VRT-043198, 2 μM Ferrostatin-1 or the DMSO vehicle. Cells were assayed by flow cytometry using SYTOX AADvanced dead cell stain. n=4 biologically independent replicates. Mean ± SD. Two-way ANOVA.

**Extended Data Fig. 3.**
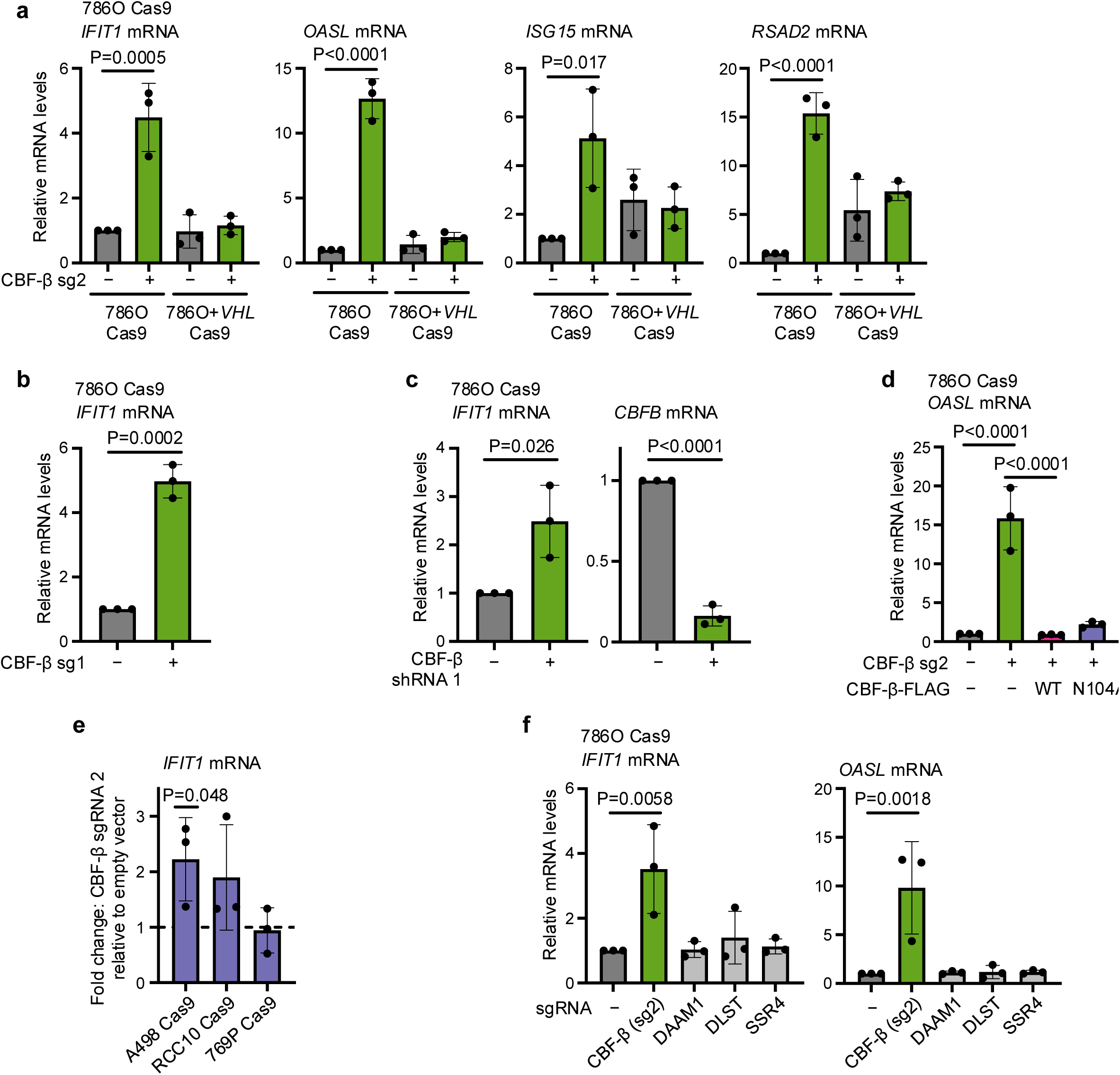
Specific induction of ISG transcription upon *CBFB* knockout (a) qPCR validation of changes in ISG expression in 786O Cas9 and 786O+*VHL* cells upon transduction with CBF-β sg2 or an empty vector control. n=3 biologically independent replicates. Mean ± SD. Two-way ANOVA. (**b,c**) qPCR analysis of changes in *IFIT1* expression in 786O Cas9 cells upon transduction with CBF-β sgRNA 1 (sg1) (**b**) or CBF-β shRNA 1 (**c**), relative to an appropriate empty vector or scrambled shRNA control. n=3 biologically independent replicates. Mean ± SD. Unpaired *t*- test. (d) qPCR analysis of *OASL* expression in 786O Cas9 cells upon transduction with CBF-β sg2 and overexpression vectors encoding CBF-β-FLAG (WT) or CBF-β-FLAG (N104A), or an empty vector control. n=3 biologically independent replicates. Mean ± SD. One-way ANOVA. (e) Expression of *IFIT1* assayed by qPCR in A498, RCC10 and 769P ccRCC cell lines upon transduction with CBF-β sg2, normalised to equivalent cells transduced with an empty vector. n=3 biologically independent replicates. Mean ± SD. Unpaired *t*-test. qPCR analysis of ISGs upon transduction with sgRNAs targeting CBF-β, DAAM1, DLST, or SSR4, relative to an empty vector-transduced control. n=3 biologically independent replicates. Mean ± SD. One-way ANOVA.

**Extended Data Fig. 4.**
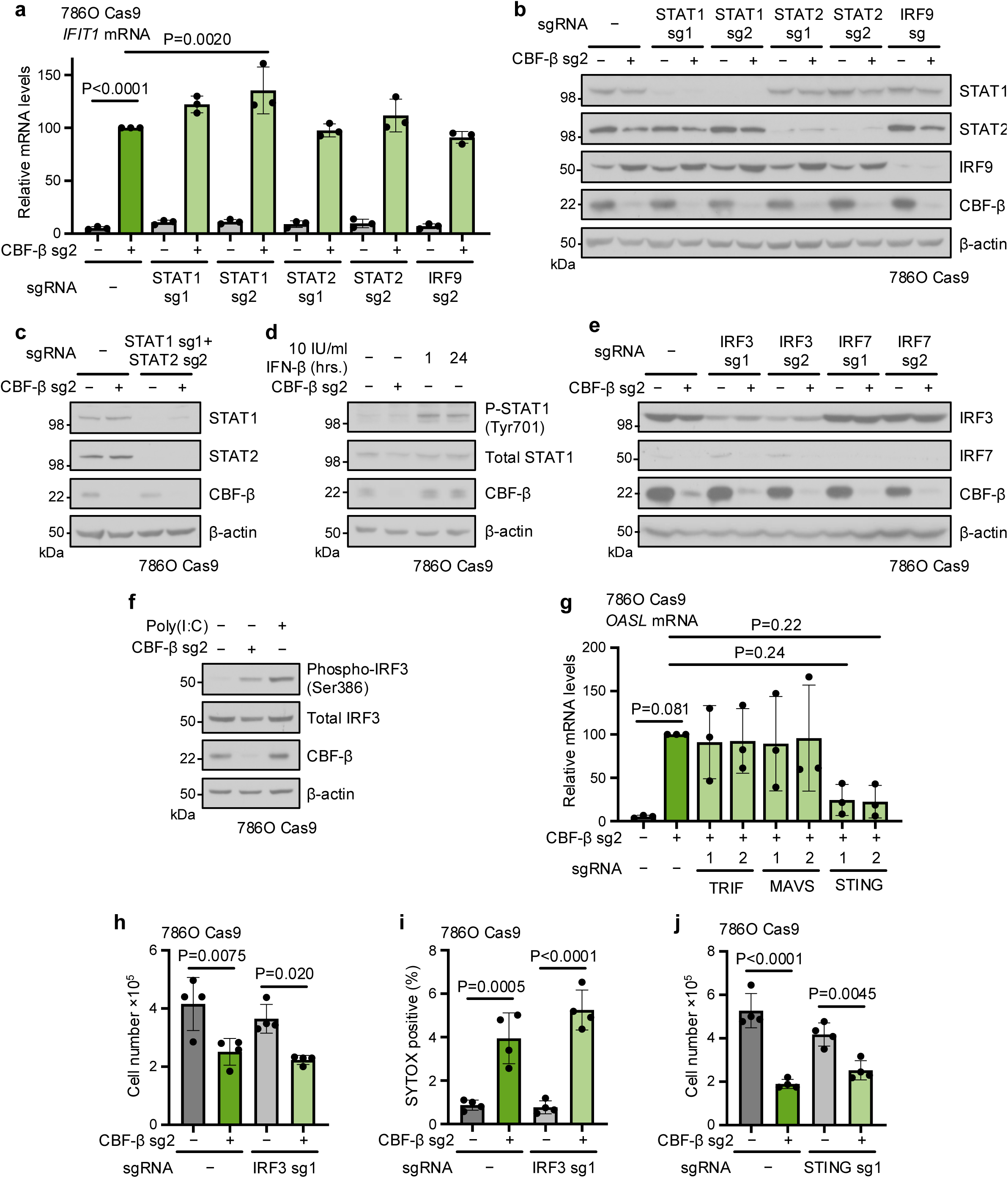
ISG transcription following *CBFB* deletion is mediated by the STING-TBK1-IRF3 axis (**a,b**) 786O Cas9 cells were transduced with CBF-β sg2 and sgRNAs targeting STAT1, STAT2 or IRF9, or an empty vector. Cells were analysed by qPCR (**a**) or immunoblot (**b**). n=3 biologically independent replicates for (**a**) and (**b**). Mean ± SD. Two-way ANOVA. (c) Immunoblot of 786O Cas9 cells transduced with CBF-β sg2, a vector encoding sgRNAs targeting both STAT1 (sg1) and STAT2 (sg2), or an empty vector control. Representative of 3 biologically independent replicates. (d) CBF-β does not affect the phosphorylation status of STAT1. Cells were transduced with CBF-β sg2 or an empty vector control. Positive controls were additionally treated with 10 IU/ml IFN-β for 1 or 24 hours. Immunoblot representative of 3 biologically independent replicates. (e) Immunoblot of experimental conditions assayed in Fig. 5c. Representative of 3 biologically independent replicates. (f) Cells were transduced with CBF-β sg2 or an empty vector control. IRF3 phosphorylation was elicited in positive control cells treated with 20 μg/ml poly(I:C) for 6 hours. Immunoblot representative of 3 biologically independent replicates. (g) qPCR analysis of *OASL* expression in conditions described in Fig. 5f. n=3 biologically independent replicates. Mean ± SD. One-way ANOVA. (**h**-**j**) 786O Cas9 cells were transduced with sgRNAs targeting CBF-β (sg2), IRF3 (sg1), or STING (sg1), or an empty vector control, and assayed by proliferation assay (**h**,**j**), or flow cytometry with SYTOX AADvanced dead cell stain (**i**). n=4 biologically independent replicates. Mean ± SD. Two-way ANOVA.

**Extended Data Fig. 5.**
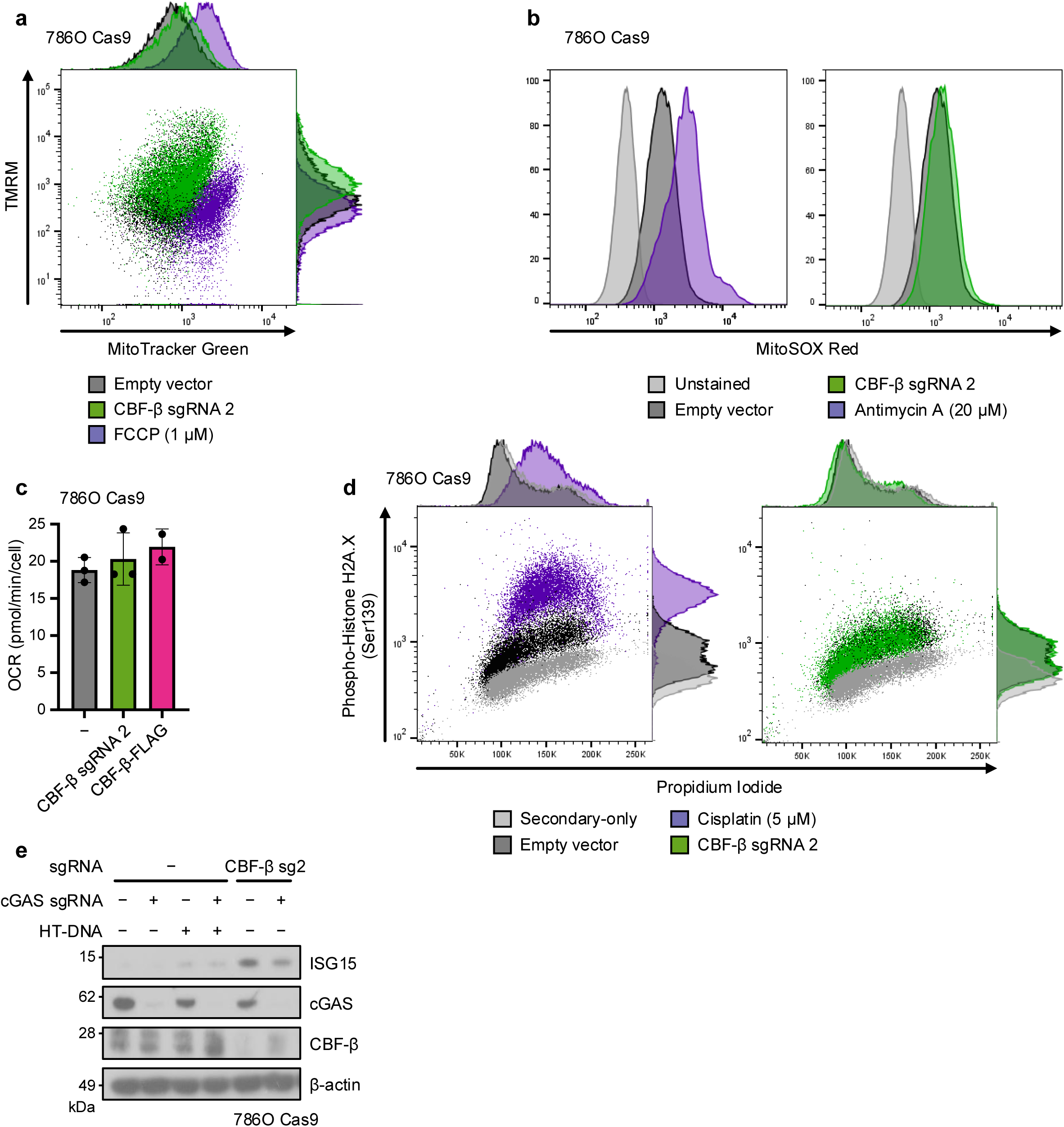
Absence of overt mitochondrial dysfunction or genomic DNA damage following CBF-β loss (**a**,**b**) 786O Cas9 cells were transduced with CBF-β sg2 or an empty vector, and analysed by flow cytometry with TMRM and MitoTracker Green FM (**a**) or MitoSOX Red (**b**) staining to assay mitochondrial membrane potential, total mitochondrial mass, and mitochondrial superoxide generation, respectively. Empty vector-transduced cells treated for 30 minutes prior to staining with inhibitors of oxidative phosphorylation, either 1 μM FCCP (**a**) or 20 μM Antimycin A (**b**), served as controls for the efficacy of staining. Representative of 3 (**a**) or 4 (**b**) biologically independent replicates. (c) 786O Cas9 cells with CBF-β depleted or overexpressed were analysed by Mito Stress Test using a Seahorse XF analyser to identify the basal level of respiration, compared to cells transduced with an empty lentiviral vector. OCR: oxygen consumption rate. n=2 (CBF-β- FLAG) or 3 (control and CBF-β sg2) biologically independent replicates. Mean ± SD. (d) CBF-β loss does not cause overt DNA damage. 786O Cas9 cells were transduced with CBF-β sg2 or an empty vector, and a population of empty vector-transduced controls were treated with 5 μM Cisplatin for 24 hours prior to analysis. Cells were stained with propidium iodide, and assessed for levels of Ser139 phosphorylation of histone 2A.X (γ-H2A.X) by intracellular antibody staining and flow cytometry. Empty vector controls were either stained fully or with the secondary antibody alone. Representative of 4 biologically independent replicates. Immunoblot of 786O Cas9 cells in the same conditions as described in Fig. 6e, following transfection with 0.5 μg/ml HT-DNA for 6 hours or having been left untransfected. Representative of 4 biologically independent replicates.

**Extended Data Fig. 6.**
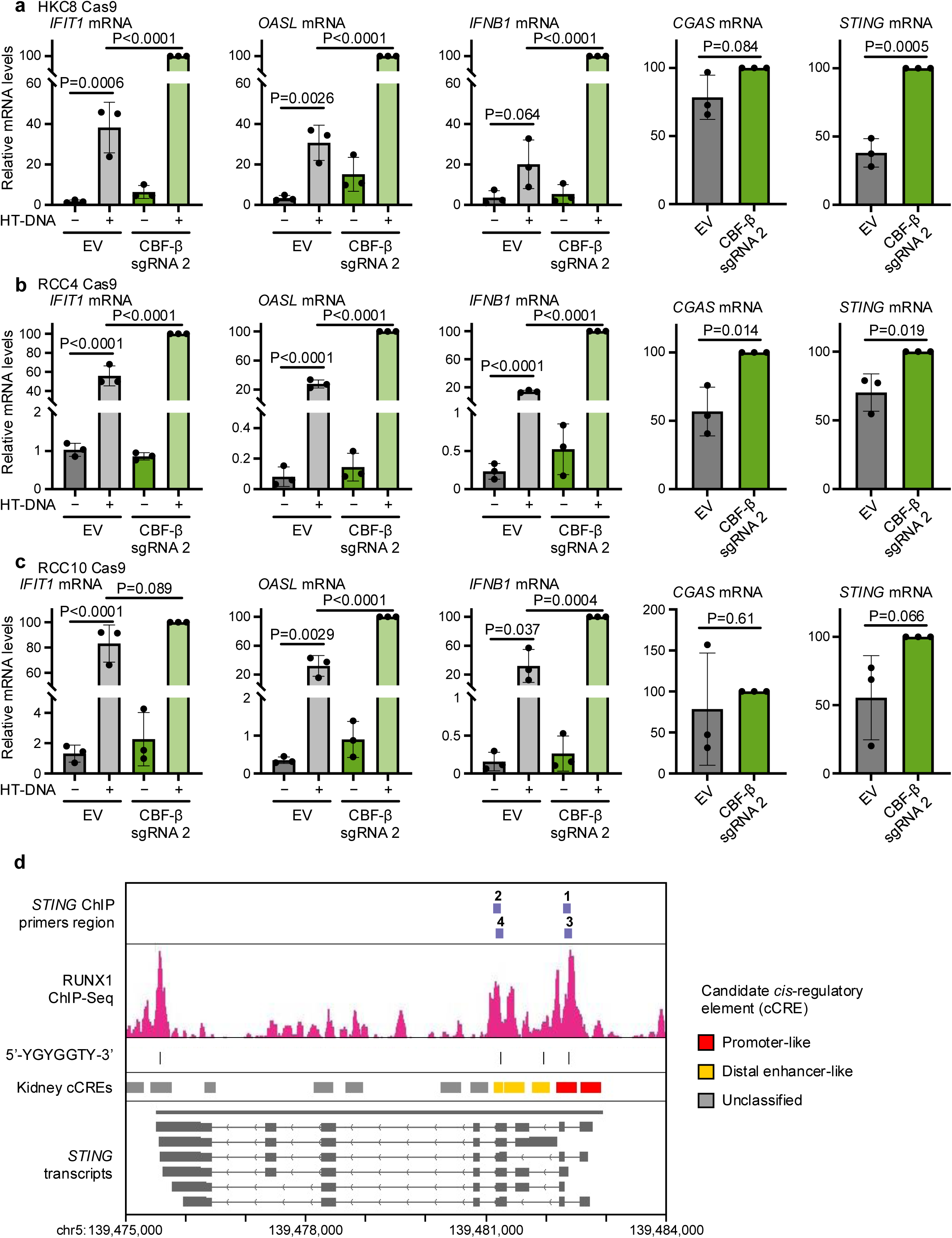
Direct transcriptional repression of STING by CBF-β/RUNX across kidney cell lines (**a-c**) The expression of *STING* and downstream ISGs is induced by *CBFB* knockout in renal proximal tubule epithelial (HKC8), and ccRCC (RCC4 and RCC10) cell lines. qPCR analysis of HKC8 Cas9 (**a**), RCC4 Cas9 (**b**), and RCC10 Cas9 (**c**) cells transduced with CBF-β sg2 or an empty vector (EV), and either transfected with 0.5 μg/ml HT-DNA for 6 hours or left untransfected. n=3 biologically independent replicates. Mean ± SD. Two-way ANOVA (*IFIT1*, *OASL* and *IFNB1*), or unpaired *t*-test (*CGAS* and *STING*). (**d**) Putative RUNX binding sites at the *STING* locus on chromosome 5. RUNX1 ChIP-Seq data from K562 cells, and human kidney candidate *cis*-regulatory element (cCRE) annotations were extracted from ENCODE files ENCFF985UVT and ENCFF657KYN, and visualised in IGV together with *STING* transcripts derived from the Ensembl genome browser. Primer pairs used for ChIP-qPCR in Fig. 6e were designed around RUNX consensus motifs (5’-YGYGGTY- 3’) in the reverse strand, which are indicated as dashes.**Extended Data** Fig. 6. Direct transcriptional repression of STING by CBF-β/RUNX across kidney cell lines (**a-c**) The expression of *STING* and downstream ISGs is induced by *CBFB* knockout in renal proximal tubule epithelial (HKC8), and ccRCC (RCC4 and RCC10) cell lines. qPCR analysis of HKC8 Cas9 (**a**), RCC4 Cas9 (**b**), and RCC10 Cas9 (**c**) cells transduced with CBF-β sg2 or an empty vector (EV), and either transfected with 0.5 μg/ml HT-DNA for 6 hours or left untransfected. n=3 biologically independent replicates. Mean ± SD. Two-way ANOVA (*IFIT1*, *OASL* and *IFNB1*), or unpaired *t*-test (*CGAS* and *STING*). (**d**) Putative RUNX binding sites at the *STING* locus on chromosome 5. RUNX1 ChIP-Seq data from K562 cells, and human kidney candidate *cis*-regulatory element (cCRE) annotations were extracted from ENCODE files ENCFF985UVT and ENCFF657KYN, and visualised in IGV together with *STING* transcripts derived from the Ensembl genome browser. Primer pairs used for ChIP-qPCR in Fig. 6e were designed around RUNX consensus motifs (5’-YGYGGTY- 3’) in the reverse strand, which are indicated as dashes.

## Supplementary information

**Supplementary Table 1.** Analysed CRISPR screen data (BAGEL2 analysis)

**Supplementary Table 2**. Analysed RNA sequencing data (gene expression analysis)

**Supplementary Table 3**. Analysed LC-MS data

**Supplementary Table 4**. Analysed RNA sequencing data (transposable elements upregulated upon CBF-β knockout)

**Supplementary Table 5.** Details of cell lines, antibodies, reagents, plasmids and software

**Supplementary Table 6**. Oligonucleotide sequences

**Supplementary Fig. 1.**
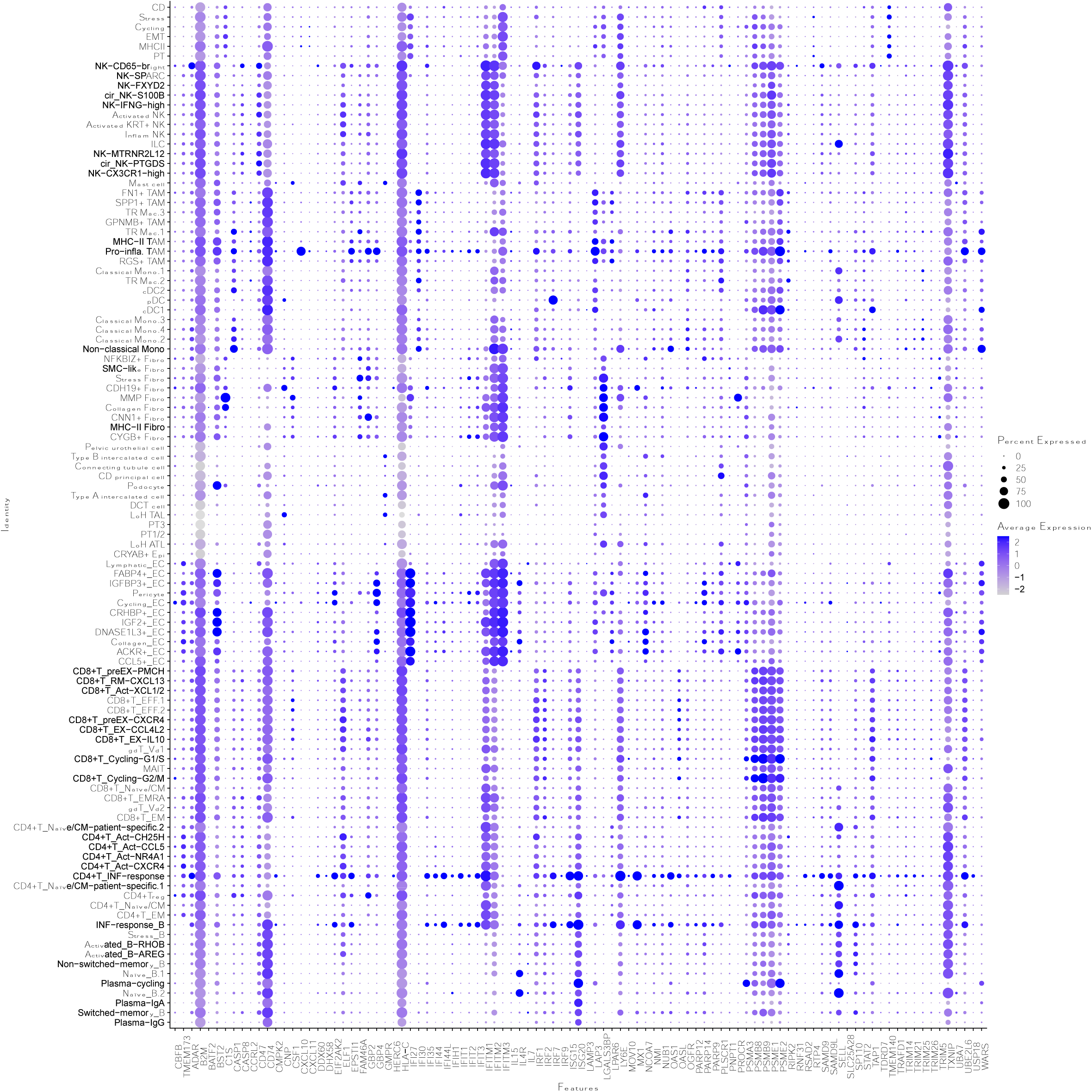
scRNA analysis of type I IFN genes in ccRCC Extended analysis of single cell transcriptomic data from patients with ccRCC. The average expression and the percentage of cells that express type I interferon genes is shown for the principal cell types within the tumour micro-environment.

**Supplementary Fig. 2.**
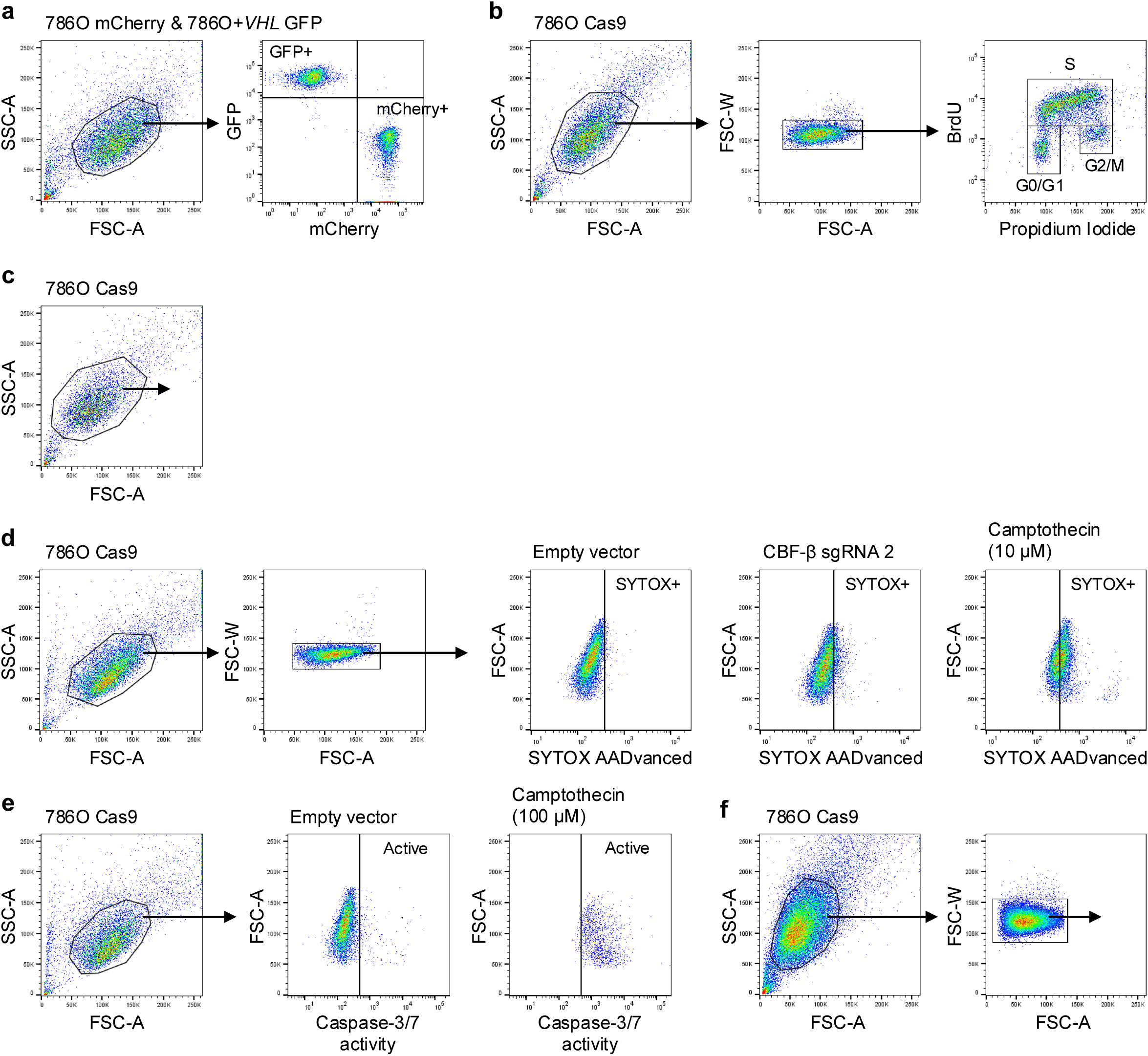
Gating strategies for flow cytometry (**a**-**f**) Representative gating strategies used in flow cytometry experiments in 786O Cas9 cells: competitive growth assay (**a**); cell cycle analysis with BrdU and PI (**b**); cell cycle analysis with PI alone (**c**); SYTOX cell death assays (**d**); Caspase-3/7 Green apoptosis assays (**e**); and staining of γ-H2A.X, and for functional mitochondrial assays (**f**). SSC-A: side scatter (area). FSC-A: forward scatter (area). FSC-W: forward scatter (width).

