## Supplementary Table 5 for "VHL synthetic lethality screens uncover CBF-β as a negative regulator of STING"

**Supplementary Table 5: Details of cell lines, antibodies, reagents, plasmids and software**

**Cell lines**

| **Cell line** | **Source** | **Identifier** |
| --- | --- | --- |
| A498 | P. Schraml lab | N/A |
| HEK293T | ATCC | Cat#CRL-3216 |
| HKC8 | P.H. Maxwell lab | N/A |
| HK2 | ATCC | Cat#CRL-2190 |
| RCC4 | P.H. Maxwell lab | N/A |
| RCC4+*VHL* | P.J. Ratcliffe lab | N/A |
| RCC10 | P.H. Maxwell lab | N/A |
| 786O | ATCC | Cat#CRL-1932 |
| 786O+*VHL* | W.G. Kaelin Jr. lab | N/A |
| 769P | P. Schraml lab | N/A |

**Antibodies**

| **Antibody** | **Source** | **Identifier** |
| --- | --- | --- |
| Rat anti-BrdU | Abcam | Cat#ab6326 |
| Rabbit anti-CBF-β | Cell Signaling Technology | Cat#62184 |
| Rabbit anti-cGAS | Cell Signaling Technology | Cat#D1D3G |
| Mouse anti-FLAG M2 | Sigma | Cat#F1804 |
| Mouse anti-GFP | Roche | Cat#11814460001 |
| Rabbit anti-H3K36me3 | Cell Signaling Technology | Cat#9763 |
| Mouse anti-HIF-1α | BD Biosciences | Cat#610959 |
| Mouse anti-HIF-1β | Cell Signaling Technology | Cat#5537S |
| Rabbit anti-IFI44 | ThermoFisher | Cat#PA5-65370 |
| Rabbit anti-IRF3 | Cell Signaling Technology | Cat#11904 |
| Rabbit anti-IRF7 | Cell Signaling Technology | Cat#13014 |
| Rabbit anti-IRF9 | Cell Signaling Technology | Cat#76684 |
| Rabbit anti-ISG15 | Santa Cruz | Cat#50366 |
| Rabbit anti-KEAP1 | Cell Signaling Technology | Cat#4334 |
| Mouse anti-MDA5 | Hertzog et al.^107^ | N/A |
| Rabbit anti-MX1 | Cell Signaling Technology | Cat#37849 |
| Rabbit anti-NRF2 | Cell Signaling Technology | Cat#8486 |
| Rabbit anti-PARP1 | Cell Signaling Technology | Cat#9542S |
| Rabbit anti-Phospho-Histone 2A.X (Ser139) | Cell Signaling Technology | Cat#9718 |
| Rabbit anti-Phospho-IRF3 (Ser386) | Cell Signaling Technology | Cat#37829 |
| Rabbit anti-Phospho-STAT1 (Tyr701) | Cell Signaling Technology | Cat#9167 |

**Reagents**

| **Reagent** | **Source** | **Identifier** |
| --- | --- | --- |
| Ampicillin | ThermoFisher | Cat#J6380706 |
| Antimycin A | Alfa Aesar | Cat#J63522 |
| Azido-PEG3-Azide | Lumiprobe | Cat#207 |
| BAY 11-7085 | MedChemExpress | Cat#HY-10257 |
| Benzonase | Sigma | Cat#E1014 |
| Blasticidin | TOKU-E | Cat#B001 |
| Bromodeoxyuridine (BrdU) | Abcam | Cat#ab142567 |
| Camptothecin | Cayman Chemical | Cat#11694 |
| CellEvent Caspase-3/7 Green detection reagent | ThermoFisher | Cat#C10423 |
| Cisplatin | Sigma | Cat#232120 |
| cOmplete EDTA-free protease inhibitor cocktail | Roche | Cat#46931590001 |
| Crystal violet | PhytoTech | Cat#C1830 |
| DMEM | Sigma | Cat#D6429 |
| DMSO | Santa Cruz | Cat#sc359032 |
| Doxycycline | Cayman Chemical | Cat#14422 |
| D-PBS | ThermoFisher | Cat#J67802.K2 |
| FCCP | Cayman Chemical | Cat#15218 |
| FCS | Sigma | Cat#F0392 |
| Ferrostatin-1 | Abcam | Cat#ab146169 |
| GSK8612 | Selleck Chemicals | Cat#S8872 |
| HBSS | Gibco | Cat#14175095 |
| Herring Testes DNA (HT-DNA) | Sigma | Cat#D6898 |
| Hoescht 33342 | Tocris Bioscience | Cat#5117/50 |
| Hygromycin | ThermoFisher | Cat#10687010 |
| IFN-β | PeproTech | Cat#300-02BC |
| Lipofectamine 2000 | Invitrogen | Cat#11668027 |
| Matrigel | Corning | Cat#354234 |
| MitoSOX Red | ThermoFisher | Cat#M36008 |
| MitoTracker Green FM | ThermoFisher | Cat#M7514 |
| Necrostatin-1s | Cayman Chemical | Cat#20924 |
| Pierce Enhanced Chemiluminescent Substrate | ThermoFisher | Cat#32209 |
| Pierce Protein G magnetic beads | ThermoFisher | Cat#88848 |
| Poly(I:C) | ApexBio | Cat#B5551 |
| Propidium iodide | Sigma | Cat#P4864 |
| Proteinase K | Thermo Scientific | Cat#EO0491 |
| Protoscript II Reverse Transcriptase | NEB | Cat#M0368X |
| PureLink DNase | ThermoFisher | Cat#12185010 |
| Purina rodent chow #5001 with 2000 ppm doxycycline | Research Diets | Cat#C11300-2000i |
| Puromycin | MP Biomedicals | Cat#200-387-8 |
| PVDF membranes | Merck | Cat#IPVH00010 |
| RNase H | Thermo Scientific | Cat#EN0202 |
| RPMI-1640 | Sigma | Cat#R8758 |
| Seahorse XF RPMI medium, pH 7.4 | Agilent | Cat#103576-100 |
| SeeBlue Plus2 | Invitrogen | Cat#LC5925 |
| SuperSignal West Dura Extended Duration Substrate | ThermoFisher | Cat#34075 |
| SuperSignal West Pico Plus Chemiluminescent Substrate | ThermoFisher | Cat#34577 |
| SYBR Green PCR Master Mix | ThermoFisher | Cat#4309155 |
| SYTOX AADvanced | ThermoFisher | Cat#S10349 |
| Thymidine | Sigma | Cat#T1895 |
| TMRM | ThermoFisher | Cat#T668 |
| TMTpro labels | ThermoFisher | Cat#A44520 |
| Trypsin/lysC mix | Promega | Cat#V5071 |
| VRT-043198 | MedChemExpress | Cat#HY-112226 |

**Plasmids**

| **Plasmid** | **Source** | **Identifier** |
| --- | --- | --- |
| Lenti‐Cas9‐2A‐Blast | Hart et al.^38^ | Cat#Addgene-73310 |
| pC.SIREN.Puro | Schaller et al.^108^ | N/A |
| pCFD5 | Port and Bullock^103^ | Cat#Addgene-73914 |
| pCMVR8.91 (Lentiviral Gag/Pol) | Demaison et al.^109^ | N/A |
| pHRSIN-pSFFV-GFP | Dupont et al.^110^ | N/A |
| pHRSIN-pSFFV-CBF-β (N104A)-FLAG-Puro | This paper | N/A |
| pHRSIN-pSFFV-CBF-β (WT)-FLAG-Puro | This paper | N/A |
| pHRSIN-pSFFV-mCherry-Puro | This paper | N/A |
| pHRSIN-pSFFV-pPGK-Puro | Demaison et al.^109^ | N/A |
| pHRSIN-SFFV-eGFP-P2A-Vif | Marelli et al.^111^ | N/A |
| pHRSIN-pSFFV-luciferase-Puro | P.J. Lehner lab | N/A |
| pKLV-U6gRNA(BbsI)-PGKblast2ABFP | P.J. Lehner lab | N/A |
| pKLV-U6gRNA(BbsI)-PGKpuro2ABFP | Koike-Yusa et al.^112^ | Cat#Addgene-50946 |
| pLCKO2-TKOv3 | Mair et al.^113^ | Cat#Addgene-125517 |
| pMD.G (Lentiviral VSVG) | Demaison et al.^109^ | N/A |
| pSpCas9(BB)-T2A-Puro | Ran et al.^101^ | Cat#Addgene-48139 |
| tet-pLKO-sgRNA-puro | Huang et al.^102^ | Cat#Addgene-104321 |

**Software**

| **Software** | **Source** | **Identifier** |
| --- | --- | --- |
| BAGEL2 | Kim and Hart^114^ | https://github.com/hart-lab/bagel |
| Cutadapt v4.4 | Martin^115^ | RRID:SCR_011841 |
| DESeq2 | Love et al.^116^ | RRID:SCR_015687 |
| ENCODE | The ENCODE Project Consortium^63^ | https://www.encodeproject.org |
| fgsea v1.20.0 | Korotkevich et al.^117^ | RRID:SCR_020938 |
| FlowJo v10 | BD Biosciences | RRID:SCR_008520 |
| HISAT2 v2.2.1 | Kim et al.^118^ | RRID:SCR_015530 |
| IGV v2.15.2 | Robinson et al.^119^ | RRID:SCR_011793 |
| limma v3.15 | Ritchie et al.^120^ | RRID:SCR_010943 |
| Peaks 11 | Bioinfor | https://www.bioinfor.com/peaks-11 |
| Prism v9.5.1 | GraphPad | RRID:SCR_002798 |
| Python v3.10 | Python Software Foundation | RRID:SCR_008394 |
| R v4.1.2 | R Core Team | RRID:SCR_001905 |
| Salmon | Patro et al.^121^ | RRID:SCR_017036 |
| STAR v2.7.10b | Dobin et al.^122^ | RRID:SCR_004463 |
| survminer v0.4.9 | Kassambara et al.^123^ | RRID:SCR_021094 |
| TEtranscripts v2.2.3 | Jin et al.^124^ | RRID:SCR_023208 |
