## Supplementary Table 6 for "VHL synthetic lethality screens uncover CBF-β as a negative regulator of STING"

**Supplementary Table 6: Oligonucleotide sequences**

**CBF-β gBlock sequence**

GTATTTTGCAGAGCTGGCGAATGCCGCGCGTCGTGCCCGACCAGAGAAGCAAGTTCGAGAACGAGGAGTTTTTTAGGAAGCTGAGCCGCGAGTGTGAGATTAAGTACACGGGCTTCAGGGACCGGCCCCACGAGGAACGCCAGGCACGCTTCCAGAACGCCTGCCGCGACGGCCGCTCGGAAATCGCTTTTGTGGCCACAGGAACCAATCTGTCTCTCCAGTTTTTTCCGGCCAGCTGGCAGGGAGAACAGCGACAAACACCTAGCCGAGAGTATGTCGACTTAGAAAGAGAAGCAGGCAAGGTATATTTGAAGGCTCCCATGATTCTGAATGGAGTCTGTGTTATCTGGAAAGGCTGGATTGATCTCCAAAGACTGGATGGTATGGGCTGTCTGGAGTTTGATGAGGAGCGAGCCCAGCAGGAGGATGCATTAGCACAACAGGCCTTTGAAGAGGCTCGGAGAAGGACACGCGAATTTGAAGATAGAGACAGGTCTCATCGGGAGGAAATGGAGGCAAGAAGACAACAAGACCCTAGTCCTGGTTCCAATTTAGGTGGTGGTGATGACCTCAAACTTCGTTAAACGCGTTGAACACTTCACAG

**Primers used to clone gene expression vectors**

| **Experiment** | **Primer** | **Sequence** |
| --- | --- | --- |
| pHRSIN-pSFFV-CBF-β (N104A)-FLAG-Puro  pHRSIN-pSFFV-CBF-β (WT)-FLAG-Puro | Fw | CAGTCCTCCGACAGACTGAGTCGCCCGGGGGGGATCC  GCCACCATGCCGCGCGTCGTGCCCGACCAGAGAAGC |
|  | Rv | CCGTCATGGTCTTTGTAGTCAGCCCGCTCGAGCGG  CCGCCCACGAAGTTTGAGGTCATCACCACCACCTAA |
|  | N104A mutation (Fw) | CTCCCATGATTCTGGCTGGAGTCTGTGTTATC |
|  | N104A mutation (Rv) | GATAACACAGACTCCAGCCAGAATCATGGGAG |
| To clone  mCherry into  pHRSIN-SFFV | Fw | CAGTCCTCCGACAGACTGAGTCGCCCGGGGGGGAT  CCGCCACCatggtgagcaagggcgaggaggataac |
|  | Rv | CTTGCATGCCTGCAGGTCGACTCTAGAGTCGCG  GCCGCTCActtgtacagctcgtccatgccgccgg |

**sgRNA sequences**

| **Target** | **sgRNA sequence** |
| --- | --- |
| CBF-β (sg1) | GAAGCTGAGCCGCGAGTGTG |
| CBF-β (sg2) | GCCTTGCAGATTAAGTACAC |
| cGAS | AAATTAAGAAGAAACATGG |
| DAAM | CAGCCGATACGTGATTTCT |
| DLST | TACACAACCTTCCTGCTGTT |
| HIF1β | CAGTCCTCCGTCTCCTCACC |
| IRF3 (sg1) | AACCAGAGGGCATAGCG |
| IRF3 (sg2) | ATCTGATTACCTTCACGGA |
| IRF7 (sg1) | ATGCTGCGGGATAACTCGG |
| IRF7 (sg2) | AAGCAGCTGCGCTACACGG |
| IRF9 | GGCTCAGCAACATCCATG |
| KEAP1 (sg1) | GTTACGGGGCACGCTCATGG |
| KEAP1 (sg2) | AGCACCGGCGAAGTGCCCTG |
| LacZ | CAGCTGGCGTAATAGCGAAG |
| MAVS (sg1) | TGTTCACAGGCATCAAGG |
| MAVS (sg2) | ACGGGAGCAGCAGAAATG |
| NRF2 (sg1) | CATTAATTCGGGATATACGT |
| NRF2 (sg2) | GGACATTGAGCAAGTTTGGG |
| RUNX1 | GCTCCGTGCTGCCTACGCAC |
| RUNX2 | GTAGGTGTGGTAGTGAGTGG |
| RUNX3 | GGACGTGCCGGATGGTACGG |
| SSR4 | TGAGGTTAGATTCTTCGACG |
| STAT1 (sg1) | ACGTTGGAGATCACCACAA |
| STAT1 (sg2) | AGGTCATGAAAACGGATGG |
| STAT2 (sg1) | TTGGCTGGCCAGAACACCG |
| STAT2 (sg2) | GGCCCAGCAAGCTCCAGG |
| STING (sg1) | CACCCCACAGTCCAATGGG |
| STING (sg2) | AAAAAGGGAATTTCAACG |
| TRIF (sg1) | AAGCTGGGCCAGGAAACTG |
| TRIF (sg2) | TGCGTGGTGGATAATGAG |

**shRNA sequences**

| **Target** | **shRNA sequence** |
| --- | --- |
| CBF-β (sh1) | TGACCTCAAACTTCGTTAATT |
| CBF-β (sh2) | GAGAAGCAGGCAAGGTATATT |
| CBF-β (sh3) | CCGCGAGTGTGAGATTAAGTA |
| Scrambled sh | GCATAATTAATATCCGCGTGT |

**qPCR primers**

| **Target** | **Forward primer** | **Reverse primer** |
| --- | --- | --- |
| *CBFB* | GCTCGGAAATCGCTTTTGTGG | CGGCTAGGTGTTTGTCGCT |
| *CGAS* | CGGGAGCTACTATGAGCACG | GCCATGTTTCTTCTTGGAAACCA |
| *GLUT1* | CCAGGGTAGCTGCTGGAGC | TGGCATGGCGGGTTGT |
| *HMOX1* | CAACATCCAGCTCTTTGAGG | GGCAGAATCTTGCACTTTG |
| *IFIT1* | ggaatacacaacctactagcc | ccaggtcaccagactcctca |
| *IFNB1* | CAGCATCTGCTGGTTGAAGA | CATTACCTGAAGGCCAAGGA |
| *ISG15* | CTCTGAGCATCCTGGTGAGGAA | AAGGTCAGCCAGAACAGGTCGT |
| *NQO1* | CCTGCCATTCTGAAAGGCTGGT | GTGGTGATGGAAAGCACTGCCT |
| *OASL* | gcggagcccatcacggtcac | agcaccaccgcaggccttga |
| *PHD3* | TCCTGCGGATATTTCCAGAGG | GGTTCCTACGATCTGACCAGAA |
| *RSAD2* | CCAGTGCAACTACAAATGCGGC | CGGTCTTGAAGAAATGGCTCTCC |
| *RUNX1* | CACTGTGATGGCTGGCAATGATG | CTCTGTGGTAGGTGGCGACTTG |
| *RUNX2* | AGCCCTCGGAGAGGTACCA | CGGAGCTCAGCAGAATAATTTTC |
| *RUNX3* | GACTGTGATGGCAGGCAATGA | CGAAGCGAAGGTCGTTGAA |
| *STING*  (qPCR) | CCTGAGTCTCAGAACAACTGCC | GGTCTTCAAGCTGCCCACAGTA |
| *STING*  (ChIP 1) | TCCACAACACTCTAGCCCTG | GGAAATACCCTCCTTCCCAGC |
| *STING*  (ChIP 2) | GGAGGATGTTCAGTGCCTGC | CTACTACTCCCTCCCAAATGCG |
| *STING*  (ChIP 3) | CCAAACCGCAGCTTTACTGG | ATCAGGGCTTTGAGGGAAGG |
| *STING*  (ChIP 4) | AAGCATCCAAGTGAAGGGCG | CCTGTTGCTGCTGTCCATCTAT |
| *VEGFA* | TACCTCCACCATGCCAAGTG | ATGATTCTGCCCTCCTCCTTC |
| *β-Actin* | CTGGGAGTGGGTGGAGGC | TCAACTGGTCTCAAGTCAGTG |

**Primers used to amplify and sequence the TKOv3 library**

| **Reaction** | **Primer** | **Sequence** |
| --- | --- | --- |
| Outer PCR | Fw | GAGGGCCTATTTCCCATGATTC |
|  | Rv | CAAACCCAGGGCTGCCTTGGAA |
| Inner PCR | Fw | AATGATACGGCGACCACCGAGATCTACACTCTCTTGTGGAAA  GGACGAGGTACCG |
|  | Rv | CAAGCAGAAGACGGCATACGAGATNNNNNNNGTGACTGGAG  TTCAGACGTGTGCTCTTCCGATCTATTTTAACTTGCTATTTCTA  GCTCTAAAAC  (NNNNNNN: Unique barcode sequence for multiplexing) |
| Illumina Sequencing | | ACACTCTCTTGTGGAAAGGACGAAACACCG |
